# MCT6 is an intestinal Lac-Phe exporter required for metformin-associated weight loss

**DOI:** 10.64898/2026.09.22.753556

**Authors:** Shuke Xiao, Alan Sheng-Hwa Tung, Jan Spaas, Sipei Fu, Xudong Chen, Tuan Kiet Trinh, Maria Dolores Moya-Garzon, Veronica L. Li, Hannah T. Cessna, Kentaro Ito, Saranya C. Reghupathy, Chao Lin, Xuchao Lyu, Steffen H. Raun, Miles D.W. Tyner, Arthur Germakovski, Jacob Tondreau, Megan Yeckley, Wei Wei, Michael R. Howitt, Mark D. Parker, Jason A. Sprowl, Stephen M. Hinshaw, Jonathan Z. Long

## Abstract

Metabolites are increasingly recognized as circulating molecules that regulate physiology, yet the mechanisms that couple intracellular production to organism-wide action remain poorly defined. Using the anorexigenic metabolite Lac-Phe as a tractable system, we identify the orphan transporter MCT6 (SLC16A5) as a physiologic intestinal Lac-Phe exporter. This mechanism controls the extent to which intracellularly synthesized Lac-Phe acquires systemic activity. MCT6 transports Lac-Phe, mediates its cellular efflux, and is required for maintaining its blood levels in mice following strong glycolytic stimuli. Both global and intestinal epithelial-specific deletion of MCT6 confers resistance to metformin-associated weight loss on a high-fat diet. Bypassing the transport defect with exogenous Lac-Phe normalizes the body weight phenotype of MCT6-KO mice. Together, these data connect MCT6 to metformin pharmacology and intestinal lactate metabolism, and more generally underscore the importance of transporter-mediated release in the conversion of an intracellular metabolic state into a circulating metabolite effector.

## Introduction

Metabolites are increasingly recognized to function as circulating molecules that communicate cellular metabolic state across tissues and regulate organismal physiology.^1^ Their levels in blood are dynamically regulated and determine the magnitude of their systemic actions. Many circulating metabolites are synthesized intracellularly, yet unlike peptide hormones or neurotransmitters, lack established pathways for vesicular storage and regulated exocytosis. Instead, extracellular release from producing cells can occur directly across the plasma membrane through the action of membrane transporters.^2^ Yet, with rare exceptions,^3–5^ the physiologically relevant exporters that mediate metabolite release into the bloodstream, and whether this step is functionally important for systemic activity, remain unknown.

Lac-Phe (*N*-lactoyl-phenylalanine) is a lactate-derived anorexigenic metabolite that provides a tractable system to define this release step and directly test its physiologic importance.^6^ Lac-Phe is synthesized intracellularly by the cytosolic enzyme CNDP2;^6,7^ intestinal epithelial CNDP2^+^ cells are a major source of the circulating pool;^8^ and Lac-Phe acts centrally on hypothalamic neurons to suppress feeding and body weight.^9^ Its circulating levels rise in response to physiologic and pharmacologic perturbations that increase glycolytic-to-oxidative ratios, including exercise,^6^ metformin treatment,^8,10^ sepsis,^11^ and mutations in the electron transport chain.^12^ Together, these features bookend the unknown intervening release step between a defined cellular source and a measurable systemic phenotype, while also providing inducible conditions in which to test its physiologic importance. Although several transporters can mediate Lac-Phe efflux in cell culture,^6,13^ none control circulating levels in vivo: SLC17A1/3 are kidney-enriched and regulate urine rather than blood Lac-Phe,^13^ while genetic ablation of ABCC5 has no effect on blood Lac-Phe.^6^ Beyond Lac-Phe, the physiologic functions of the broader *N*-lactoyl amino acid family are still largely unexplored.

MCT6 (SLC16A5) is an orphan member of the monocarboxylate transporter (MCT) family.^14^ This is a large family of transporters involved in the translocation of carboxylate-containing metabolites.^15^ The most well-studied members of this family, MCT1 (SLC16A1) and MCT4 (SLC16A3), are lactate transporters.^16,17^ MCT6 shares 30-40% sequence identity with MCT1 and MCT4, but does not itself transport lactate.^18^ The endogenous physiologic substrates and functions of MCT6 remain unknown, and no metabolic phenotypes have been reported for MCT6-KO mice.^19,20^

Here we show that the orphan transporter MCT6 (SLC16A5) is a physiologic Lac-Phe exporter. MCT6 transports Lac-Phe in vitro, mediates Lac-Phe efflux in cells, and is required for normal circulating Lac-Phe levels in vivo, particularly after strong glycolytic stimuli. Under standard chow or high fat diet feeding conditions, MCT6-KO mice exhibit normal body weight gain; however, following a metformin challenge, high fat diet-fed MCT6-KO mice exhibit resistance to metformin-associated weight loss. Bypass of the export step by administration of exogenous Lac-Phe to metformin-treated MCT6-KO mice restores the body weight to that of metformin-treated wild-type mice. Beyond Lac-Phe, we identify other *N*-lactoyl amino acids, particularly Lac-Tyr (*N*-lactoyl-tyrosine) and Lac-Trp (*N*-lactoyl-tryptophan) as additional physiologic MCT6 substrates. However, Lac-Tyr and Lac-Trp have no effect on feeding or body weight. Intestinal epithelial-specific MCT6-KO mice also exhibit reduced metformin-inducible Lac-Phe in blood and resistance to metformin-associated weight loss, localizing the phenotype to the gut. These data deorphanize MCT6 as a Lac-Phe transporter that links gut lactate metabolism to metformin pharmacology and energy balance. More broadly, our study defines cellular export as a critical link that couples the intracellular production to the systemic activity of a metabolite effector.

## Results

### MCT6 is sufficient to drive Lac-Phe efflux in cell culture

To identify new Lac-Phe transporters that might be responsible for its efflux to the circulation, we performed a co-expression analysis between mRNA levels of candidate transporters and *Cndp2* using single cell data from the gut. Conceptually, the premise of this analysis was that transporters relevant for Lac-Phe efflux might be expected to be enriched in the same cells as the Lac-Phe biosynthetic enzyme *Cndp*2 (i.e., intestinal epithelial cells). Using data from Han et al.,^21^ we identified six cell clusters with high *Cndp2* expression and used these cells to calculate the Pearson correlation coefficient between *Cndp2* expression and that of 220 candidate transporters (**Fig. 1A** and **Table S1**). A total of 35 candidate transporters exhibited significant co-expression with *Cndp2*. We confirmed that the majority (86%, 30 of 35) of these candidate transporters were also ranked amongst the top co-expressed with *Cndp2* mRNA in an independent scRNA-seq dataset from Haber et al.^22^ (**Table S1**). For functional testing, we prioritized those co-expressed transporters with high normalized expression in the gut (**Table S1** and **Fig. 1A**). We transfected FLAG-tagged candidate transporters to HEK293T cells and measured Lac-Phe levels in media and cell lysates (**Fig. 1B**). As a control, we also included SLC17A3, a previously identified renal Lac-Phe transporter (**Fig. 1B**). Transfection of MCT6 (SLC16A5) increased media Lac-Phe levels by ∼3-fold and concurrently reduced intracellular Lac-Phe levels by ∼50% (**Fig. 1C,D**). No other candidate tested increased media Lac-Phe levels (**Fig. 1C,D**). The magnitude of the MCT6-dependent changes in media and intracellular Lac-Phe levels was comparable to that of SLC17A3 (**Fig. S1A,B**). MCT6 transfection did not change free lactate or free phenylalanine, either extracellularly or intracellularly (**Fig. S1C-F**), nor did it change CNDP2 protein levels (**Fig. S1G**), indicating that the increase in media Lac-Phe is not due to broader changes in lactate or amino acid metabolism or to Lac-Phe synthesis. Throughout this work, we use the current nomenclature MCT6/SLC16A5 (**Fig. S1H**).

**Figure 1.**
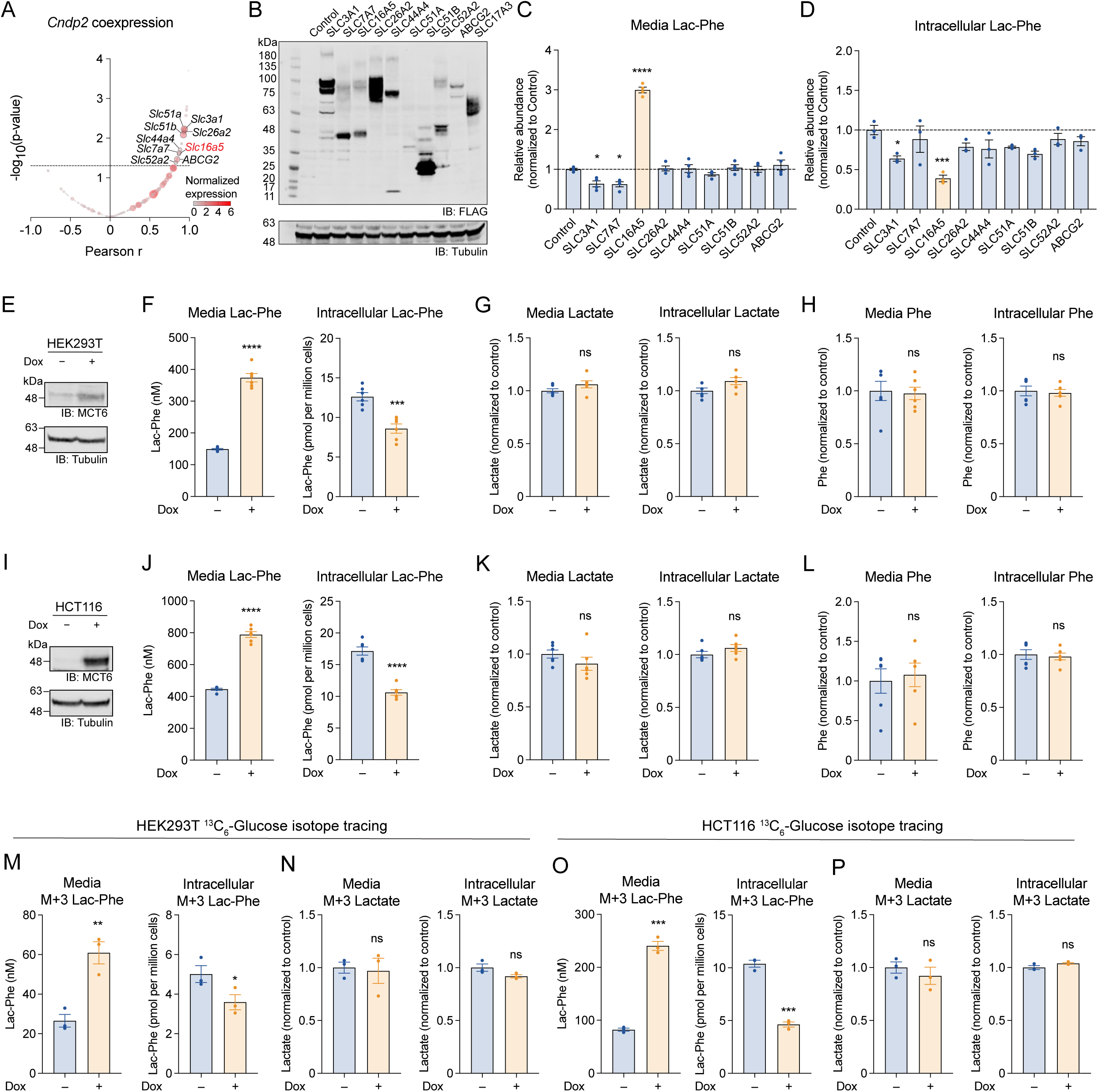
MCT6 regulates intracellular and extracellular Lac-Phe levels in cells. (A) Pearson correlation coefficient (r) and p-value (uncorrected) between mRNA levels of *Cndp2* and 220 candidate transporter genes (analyzed from Han et al. 2018). (B) Anti-FLAG (top) and anti-tubulin (bottom) blotting of HEK293T cells transfected with top CNDP2 co-expressed transporters. Control plasmid expresses mCherry. SLC17A3 was included as a positive control. (C) Lac-Phe levels in conditioned medium of transfected HEK293T cells. N = 4/group. (D) Intracellular Lac-Phe levels in transfected HEK293T cells. N = 3/group. One well was used for the Western blot in panel (B). (E) Anti-MCT6 (top) or anti-tubulin (bottom) blotting of cell lysates of doxycycline-inducible MCT6-expressing HEK293T cells. (F) Levels of Lac-Phe in conditioned medium (left) or cell lysates (right) of HEK293T cells with or without doxycycline-inducible MCT6 expression. N = 6/group. (G) Levels of lactate in conditioned medium (left) or cell lysates (right) of HEK293T cells with or without doxycycline-inducible MCT6 expression. N = 6/group. (H) Levels of phenylalanine in conditioned medium (left) or cell lysates (right) of HEK293T cells with or without doxycycline-inducible MCT6 expression. N = 6/group. (I) Anti-MCT6 (top) or anti-tubulin (bottom) blotting of cell lysates of doxycycline-inducible MCT6-expressing HCT116 cells. (J) Levels of Lac-Phe in conditioned medium (left) or cell lysates (right) of HCT116 cells with or without doxycycline-inducible MCT6 expression. N = 6/group. (K) Levels of lactate in conditioned medium (left) or cell lysates (right) of HCT116 cells with or without doxycycline-inducible MCT6 expression. N = 6/group. (L) Levels of phenylalanine in conditioned medium (left) or cell lysates (right) of HCT116 cells with or without doxycycline-inducible MCT6 expression. N = 6/group. (M) Levels of (m+3) Lac-Phe in conditioned medium (left) or cell lysates (right) of HEK293T cells with or without doxycycline-inducible MCT6 expression. N = 3/group. (N) Levels of (m+3) lactate in conditioned medium (left) or cell lysates (right) of HEK293T cells with or without doxycycline-inducible MCT6 expression. N = 3/group. (O) Levels of (m+3) Lac-Phe in conditioned medium (left) or cell lysates (right) of HCT116 cells with or without doxycycline-inducible MCT6 expression. N = 3/group. (P) Levels of (m+3) lactate in conditioned medium (left) or cell lysates (right) of HCT116 cells with or without doxycycline-inducible MCT6 expression. N = 3/group. Data are shown as mean ± SEM. * *p* < 0.05, ** *p* < 0.01, *** *p* < 0.001, **** *p* < 0.0001. In (C,D), *p*-values were calculated from one-way ANOVA with Dunnett multiple comparisons tests. In (F,G,H,J,K,L), *p*-values were calculated from two-tailed unpaired t-test with Welch’s correction. In (M,N,O,P), *p*-values were calculated from one-tailed unpaired t-test with Welch’s correction.

To further substantiate the role of Lac-Phe as an endogenous MCT6 substrate in cells, we generated stable, doxycycline (dox)-inducible, epitope (Twin-Strep and HA)-tagged MCT6-overexpressing human kidney (HEK293T) and human colon (HCT116) cell lines (**Fig. 1E-L**). In both cell lines, we detected moderate basal expression of endogenous MCT6 as measured by qPCR (Ct ∼20-26, **Fig. S1I,J**). Dox-inducible MCT6 overexpression was confirmed by anti-MCT6 Western blotting (**Fig. 1E,I**). In both cell lines, MCT6 expression once again increased media Lac-Phe while reducing intracellular Lac-Phe levels (**Fig. 1F,J**). As expected, neither lactate nor phenylalanine changed in media or lysates following MCT6 overexpression (**Fig. 1G,H** and **Fig. 1K,L**). In addition, neither CNDP2 protein levels (**Fig. S1K**) nor Lac-Phe synthesis activity (**Fig. S1L,M**) were changed in dox-inducible MCT6-overexpressing cells.

Lastly, we performed tracing experiments using heavy ^13^C_6_-glucose isotope (m+6) and monitored incorporation of the (m+3) lactate unit into Lac-Phe. In both dox-inducible HEK293T and HCT116 overexpressing lines, media (m+3) Lac-Phe was increased compared to control lines, with a reciprocal reduction in intracellular (m+3) Lac-Phe (**Fig. 1M,O**). By contrast, (m+3) lactate in both media and cells was identical between control and MCT6-overexpressing cells (**Fig. 1N,P**), demonstrating that the observed differences are specific to Lac-Phe rather than differences in (m+6) glucose or (m+3) lactate loading. We conclude that overexpression of MCT6 is sufficient to increase Lac-Phe efflux in cell culture.

### MCT6 is required for Lac-Phe efflux in cells

We next used CRISPR/Cas9 and independent guide RNAs to generate two pooled knockout lines in HCT116 cells (KO1 and KO2). The two pooled knockout lines were validated by gDNA sequencing and exhibited 75% and 93% indel rates, respectively (**Fig. S1N**). Media Lac-Phe was reduced in both MCT6-KO1 and KO2 cell lines (**Fig. 2A**). Intracellular Lac-Phe levels were also elevated in KO2, but not KO1 cells (**Fig. 2A**). Lactate and phenylalanine were not consistently changed in media or lysates from MCT6-KO cells (**Fig. 2B,C**). Lac-Phe synthesis activity and CNDP2 protein levels were also not different in MCT6-KO cells (**Fig. S1O,P**).

**Figure 2.**
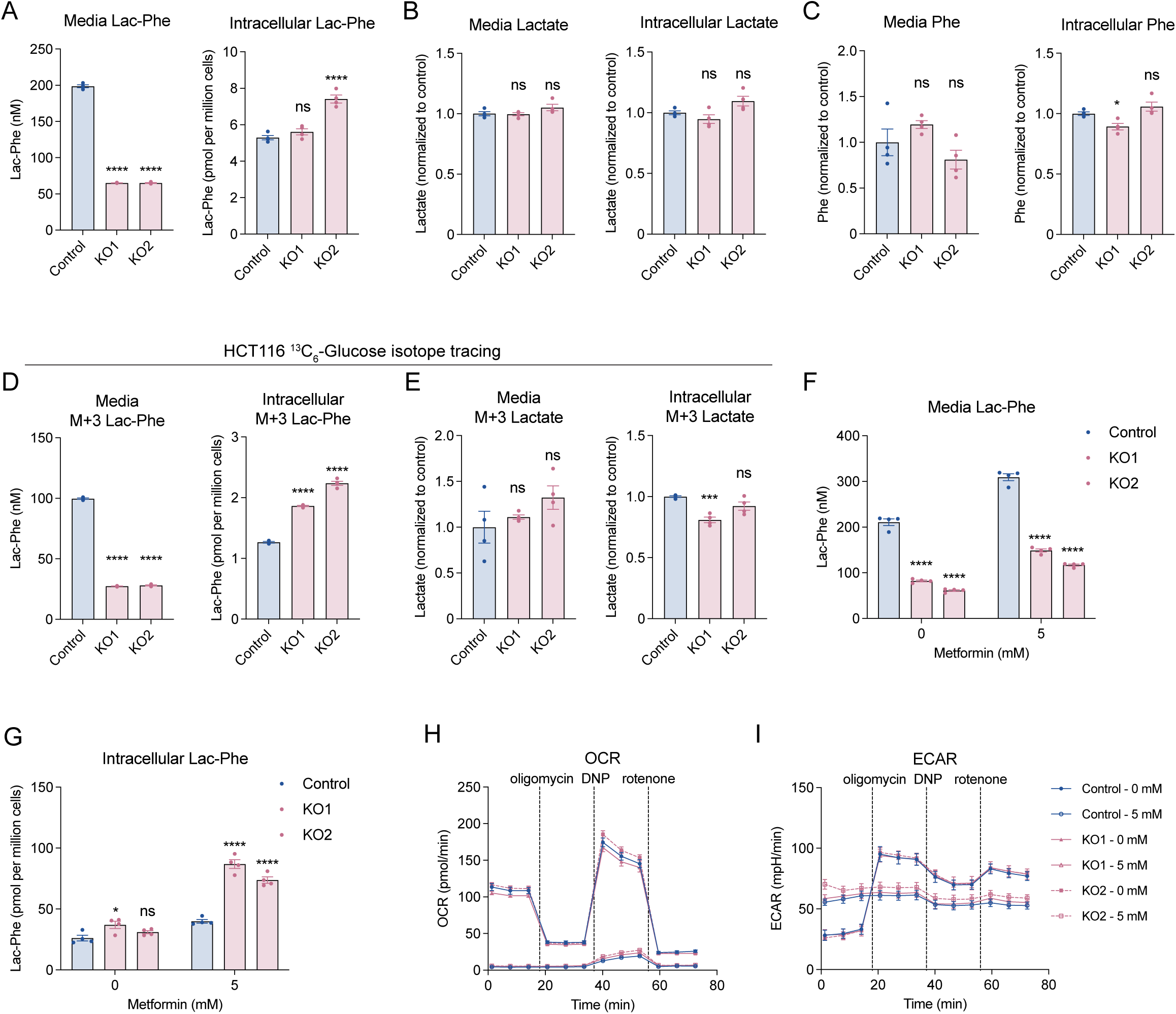
MCT6 is required for Lac-Phe efflux in cells. (A) Levels of Lac-Phe in conditioned medium (left) or cell lysates (right) of control and MCT6-KO1, KO2 HCT116 cells. N = 4/group. (B) Levels of lactate in conditioned medium (left) or cell lysates (right) of control and MCT6-KO1, KO2 HCT116 cells. N = 4/group. (C) Levels of phenylalanine in conditioned medium (left) or cell lysates (right) of control and MCT6-KO1, KO2 HCT116 cells. N = 4/group. (D) Levels of (m+3) Lac-Phe in conditioned medium (left) or cell lysates (right) of control and MCT6-KO1, KO2 HCT116 cells. N = 4/group. (E) Levels of (m+3) lactate in conditioned medium (left) or cell lysates (right) of control and MCT6-KO1, KO2 HCT116 cells. N = 4/group. (F) Levels of Lac-Phe in conditioned medium of control and MCT6- KO1, KO2 HCT116 cells after 0 or 5 mM metformin treatment. N = 4/group. (G) Levels of Lac-Phe in cell lysates of control and MCT6-KO1, KO2 HCT116 cells after 0 or 5 mM metformin treatment. N = 4/group. (H) Oxygen consumption rate (OCR) of control and MCT6-KO1, KO2 HCT116 cells following 0 or 5 mM metformin treatment. N = 9/group. (I) Extracellular acidification rate (ECAR) of control and MCT6-KO1, KO2 HCT116 cells following 0 or 5 mM metformin treatment. N = 9/group. Data are shown as mean ± SEM. * *p* < 0.05, *** *p* < 0.001, **** *p* < 0.0001. In (A-E), *p*-values were calculated from one-way ANOVA with Dunnett multiple comparisons tests. In (F,G), *p*-values were calculated from two-way ANOVA with Dunnett multiple comparisons tests.

We performed similar experiments with ^13^C_6_-glucose to measure production and efflux of newly synthesized Lac-Phe in control and MCT6-KO cells. A ∼75% reduction in media (m+3) Lac-Phe was observed in both MCT6-KO lines, with a reciprocal ∼50% increase in intracellular (m+3) Lac-Phe (**Fig. 2D**), while levels of (m+3) lactate itself were not consistently different between the two MCT6-KO lines and control cells (**Fig. 2E**).

Metformin increases Lac-Phe levels via inhibition of complex I and the subsequent increases in intracellular glycolytic (and lactate) flux.^23^ We therefore treated control and MCT6-KO cells with metformin and measured Lac-Phe in media and cells. Media Lac-Phe was consistently reduced in both MCT6-KO cell lines relative to control cells at both baseline and 5 mM metformin (**Fig. 2F**), while intracellular Lac-Phe was clearly increased in MCT6-KO cells after metformin (**Fig. 2G**). Importantly, mitochondrial function was equivalent between WT and MCT6-KO cells both at baseline and in response to metformin: basal OCR did not differ, and metformin suppressed OCR equally in both genotypes (**Fig. 2H**). Extracellular acidification rate (ECAR) was likewise comparable between WT and MCT6-KO cells under both basal and metformin treated conditions (**Fig. 2I**). We conclude that MCT6 is required for both basal and metformin-stimulated Lac-Phe efflux in cell culture.

### Transport activity, structural modeling, and mutagenesis of MCT6

To directly characterize MCT6 transport of Lac-Phe in vitro, we first reconstituted MCT6 transport activity in *Xenopus laevis* oocytes. Oocytes were injected with cRNA encoding mouse MCT6 and expression was confirmed by anti-MCT6 Western blotting (**Fig. 3A**). We observed time- and concentration-dependent MCT6-specific Lac-Phe transport that fit Michaelis-Menten kinetics with a *K*_m_ of 2.7 mM and a *V*_max_ of 62.6 pmol/min/oocyte (**Fig. 3B,C**). In parallel, we also developed a mammalian cell-based transport assay using MCT6-KO HCT116 cells reconstituted with dox-inducible MCT6 overexpression (versus non-dox treated controls) and a deuterated Lac-Phe substrate (D_3_-Lac-Phe). Once again, we observed time- and concentration-dependent MCT6-specific D_3_-Lac-Phe transport in this cell-based assay (**Fig. S2A,B**). Using the cell-based assay, we found that bumetanide, a xenobiotic and previously reported MCT6 substrate, inhibited MCT6 transport activity in a concentration-dependent manner (**Fig. S2C**). In addition, replacement of extracellular Na⁺ with NMDG^+^ (N-methyl-D-glucamine), Cl⁻ with gluconate, or NaCl with mannitol did not significantly alter MCT6 transport activity, indicating that MCT6-mediated transport did not require extracellular Na⁺ or Cl⁻ under these conditions (**Fig. S2D**). MCT6 transport activity was also stimulated by ∼4-fold at acidic versus neutral or basic pH (**Fig. S2E**). When the membrane potential and transmembrane proton gradient were jointly dissipated using high K⁺, valinomycin, and nigericin, MCT6 transport activity increased at pH 7.4 and no longer differed between pH 7.4 and pH 6.0 (**Fig. S2F**). We conclude that MCT6-mediated Lac-Phe transport is Na⁺/Cl⁻-independent, pH-sensitive, and modulated by the membrane electrochemical state.

**Figure 3.**
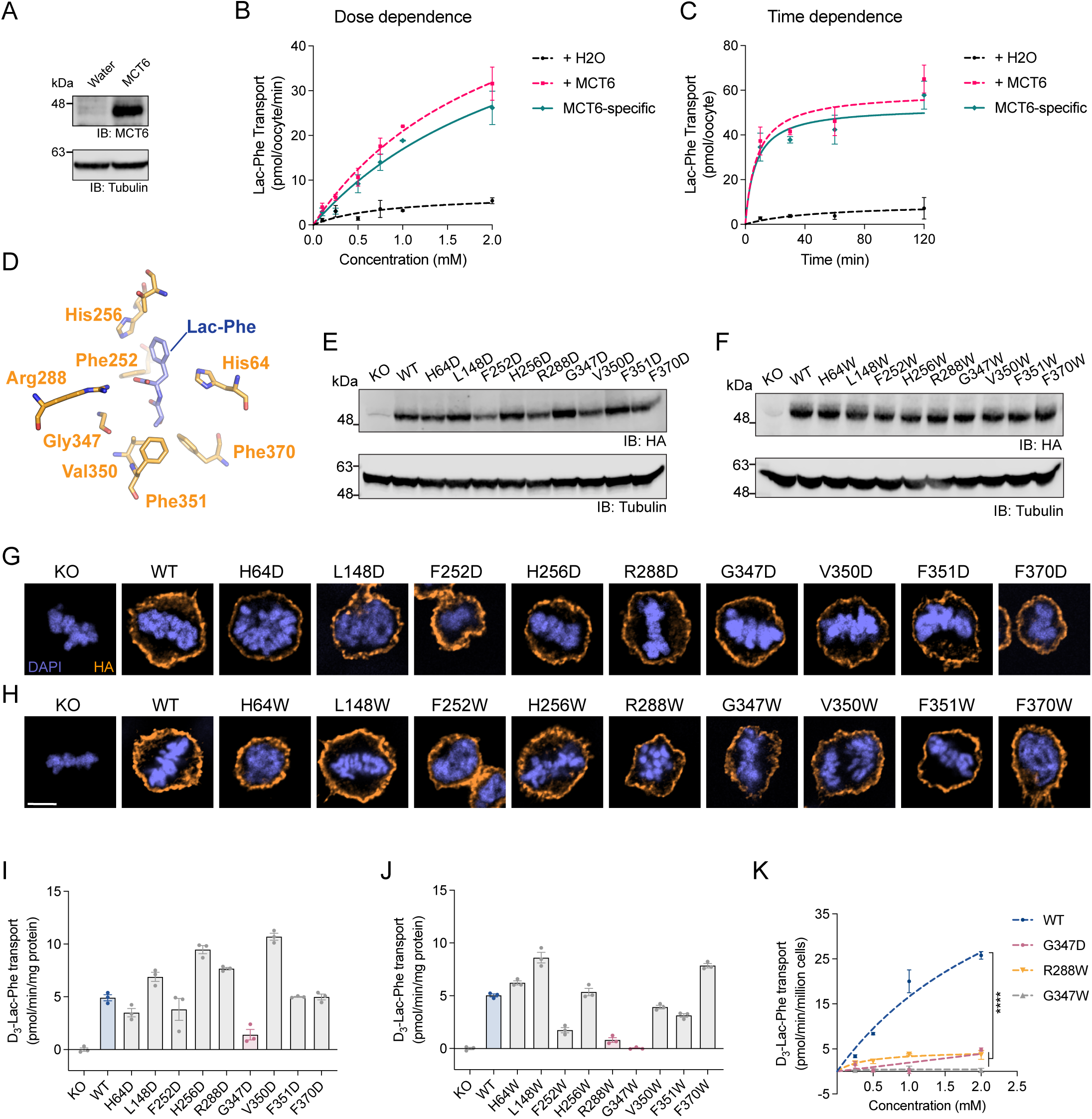
In vitro transport activity and mutagenesis of MCT6. (A) Anti-MCT6 (top) or anti-tubulin (bottom) blotting of cell lysates of *X. laevis* oocytes injected with water or MCT6 cRNA. (B,C) Lac-Phe transport activity of *X. laevis* oocytes injected with water (black line) or MCT6 cRNA (pink line), as well as MCT6-specific Lac-Phe transport activity (teal line), with different Lac-Phe concentrations after a 10 minute incubation (B) or with different incubation times and 1 mM Lac-Phe (C), N = 2-3/group. (D) Molecular docking of mMCT6 and Lac-Phe. MCT6 residues (orange) within 4Å of Lac-Phe (blue). (E,F) Anti-HA (top) or anti-tubulin (bottom) blotting of cell lysates of HCT116 MCT6-KO cells, or MCT6-KO cells expressing WT or various MCT6 mutants. (G,H) Immunofluorescence staining using DAPI and anti-HA antibodies of HCT116 MCT6-KO cells, or MCT6-KO cells expressing WT or various MCT6 mutants. Scale bar is 5 µm. (I,J) MCT6-specific D_3_-Lac-Phe transport activity of HCT116 MCT6-KO cells, or MCT6-KO cells expressing WT or various MCT6 mutants. N = 3/group. (K) MCT6-specific D_3_-Lac-Phe transport activity of WT, G347D, R288W, and G347W MCT6 mutants. N = 2-3/group. For (B,C) and (I-K), MCT6-specific transport activity was determined by subtraction of the background activity in water-injected oocytes (B,C) or MCT6-KO cells (I-K). Data are shown as mean ± SEM. **** *p* < 0.0001. In (K), *p*-values were calculated from mixed-effects analysis with Tukey’s multiple comparisons tests.

We next sought to identify MCT6 point mutants with reduced Lac-Phe transport activity. We used DiffDock to dock Lac-Phe into an AlphaFold2-modeled MCT6 protein,^24,25^ which adopted an inward-open conformation by analogy to crystal structures of other SLC family transporters (**Fig. S2G**). The docked poses were highly concordant, with the top-ranked poses all placing the benzyl ring in a hydrophobic pocket at the core of the channel (**Fig. S2H**). Based on this modeling, we identified the following candidate residues with side chains within 4 Å of Lac-Phe: His64, Leu148, Phe252, His256, Arg288, Gly347, Val350, Phe351, and Phe370 (**Fig. 3D**). We generated two sets of mutants in which these residues were individually mutated to aspartate (D) or tryptophan (W). Mutants or WT MCT6 were stably introduced into MCT6-KO HCT116 cells by viral transduction. All mutants exhibited comparable protein expression by Western blot (**Fig. 3E,F**) and plasma membrane localization by immunofluorescence microscopy (**Fig. 3G,H**). Using the cell-based D_3_-Lac-Phe transport assay, we found that most mutants retained (or in some cases increased) transport activity, whereas mutation of Gly347 strongly reduced activity in both the aspartate and tryptophan series (G347D, G347W, **Fig. 3I,J**). R288W also strongly reduced transport activity (**Fig. 3J**). The reduced transport activity of G347D, G347W, and R288W mutants were observed at all concentrations of D_3_-Lac-Phe tested (**Fig. 3K**). These three mutants also exhibited reduced bumetanide transport activity (**Fig. S2I**). These data define Gly347 and Arg288 as critical determinants of MCT6-dependent transport activity.

Lastly, to compare the predicted structural and physicochemical properties of substrate-binding cavities in MCT6, SLC17A1, SLC17A3, and ABCC5, we analyzed structural models of all four transporters using SiteMap^26^ (**Fig. S2J**). The top-ranked site in each model had a SiteScore greater than 1.10 and a Dscore greater than 1.13. MCT6 had the highest SiteScore and Dscore among the four transporters, although the values were similar across the set. The predicted MCT6 and SLC17A3 cavities had comparably high hydrophobic character, with Phobic scores of 1.562 and 1.554 and hydrophobic-to-hydrophilic balance values of 2.551 and 2.499, respectively. Both cavities were less hydrophilic than the corresponding sites in SLC17A1 and ABCC5, which had balance values of 1.486 and 1.297, respectively.

### Reduced circulating Lac-Phe levels and blunted metformin-associated weight loss in global MCT6-KO mice

We next examined the contribution of MCT6 to circulating Lac-Phe in vivo using global MCT6-KO mice and littermate wild-type (WT) controls.^20^ We confirmed loss of MCT6 protein by anti-MCT6 Western blotting of gut tissues from MCT6-KO mice, while CNDP2 protein levels were unchanged (**Fig. 4A**). In MCT6-KO mice, Lac-Phe levels in blood were reduced by ∼50% at both the basal state and following a single oral dose of metformin (300 mg/kg, p.o., **Fig. 4B**). We confirmed metformin levels in blood after dosing were equivalent between the two genotypes (**Fig. S3A**). Tissue Lac-Phe levels in gut, muscle, liver, and brain were not significantly different between genotypes at baseline or after metformin (**Fig. S3B-E**). Therefore, MCT6 is a physiologic regulator of circulating Lac-Phe levels in vivo.

**Figure 4.**
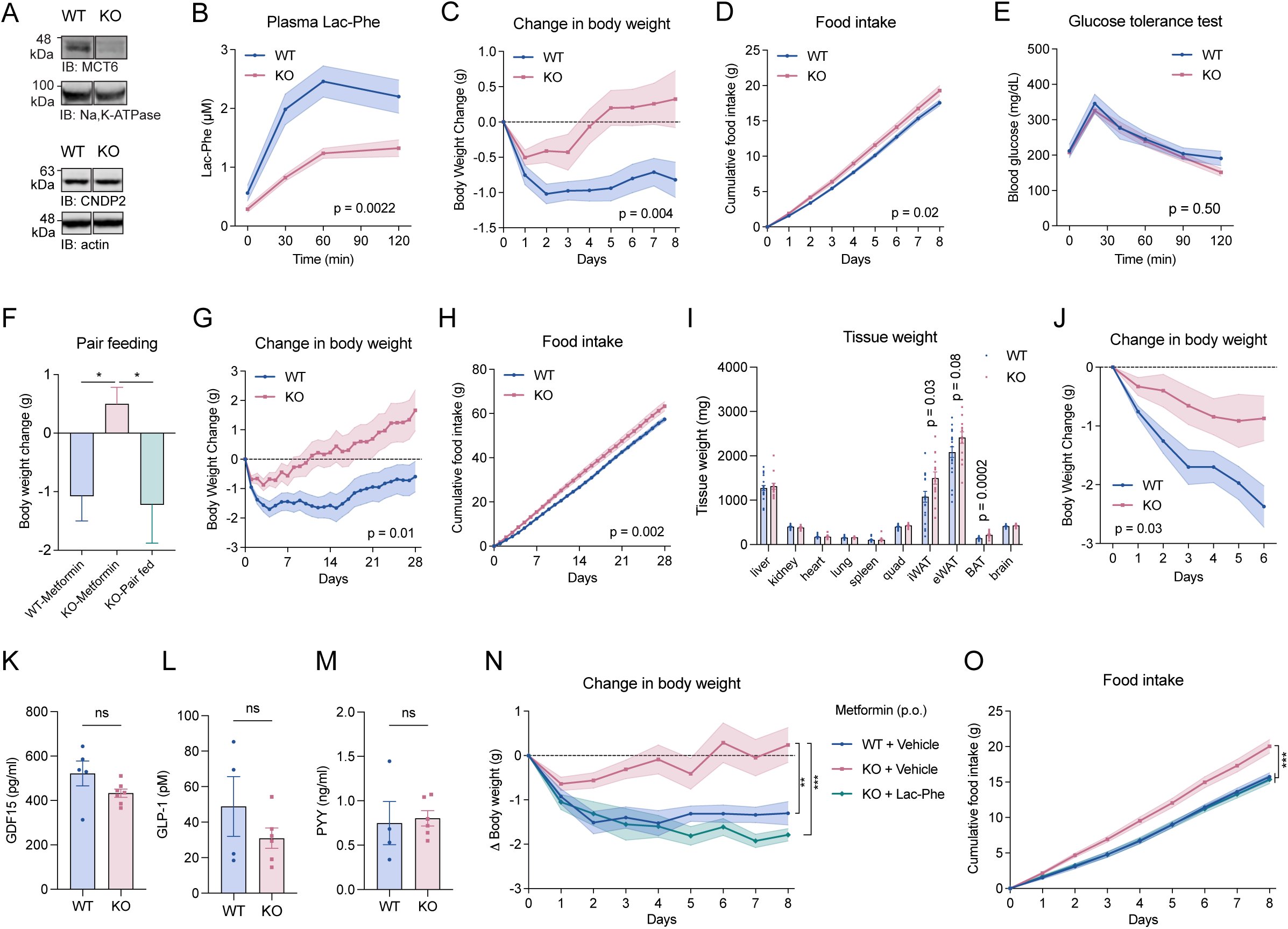
MCT6 is required for metformin-associated weight loss in mice. (A) Anti-MCT6 or anti-sodium potassium ATPase blotting (top), and anti-CNDP2 or anti-beta-actin blotting (bottom) of intestine lysates of WT and MCT6-KO mice. (B) Plasma Lac-Phe levels at the indicated time point from 8-12 week-old male WT and MCT6-KO mice before (time 0) and after metformin administration (300mg/kg, p.o.). N = 7-8/group. (C,D) Cumulative change in body weight (C) and food intake (D) in 10-14 week-old male WT and MCT6-KO DIO mice following chronic metformin treatment (300 mg/kg daily, p.o.). N=17/group. Starting body weights were WT: 35.2 ± 1.1 g, KO: 34.6 ± 1.5 g (mean ± SEM). (E) Glucose levels of WT and MCT6-KO DIO male mice (12-14 week-old) during glucose tolerance test. Metformin (300 mg/kg, p.o.) was administered 30 minutes before glucose administration (1 g/kg, i.p.). N = 7 for WT, N = 11 for MCT6-KO. (F) Body weight change of single-housed 12-19 week-old male DIO mice in three groups: WT (WT-Metformin) and MCT6-KO (KO-Metformin) mice after 14 days treatment with metformin, MCT6-KO pair-fed with the average food intake of WT-Metformin mice. N = 8-9/group. Starting body weights were WT-Metformin: 36.8 ± 1.5 g, KO-Metformin: 36.1 ± 1.9 g, and KO-Pair-fed: 37.8 ± 2.7 g (mean ± SEM). (G,H) Cumulative change in body weight (G) and food intake (H) in 17-18 week-old male WT and MCT6-KO DIO mice following chronic metformin treatment (300 mg/kg daily, p.o.). N = 20 for WT, N = 14 for KO. Starting body weights were WT: 37.1 ± 0.7 g, KO: 36.7 ± 1.4 g (mean ± SEM). (I) Tissue weights of male WT and MCT6-KO DIO mice after chronic metformin treatment for 28 days. N = 20 for WT, N = 14 for KO. (J) Cumulative change in body weight in 10-14 week old male WT and MCT6-KO DIO mice following chronic metformin treatment (600 mg/kg daily, p.o). N = 7/group. Starting body weights were WT: 42.0 ± 1.4 g, KO: 40.0 ± 1.7 g (mean ± SEM). (K-M) Plasma GDF15 (K) or GLP-1 (L) or PYY (M) levels in 12-14 week old male WT and MCT6-KO DIO mice 4 hours after a single administration of metformin (300 mg/kg, p.o.). N = 4-7/group. (N,O) Cumulative change in body weight (N) and food intake (O) in 15-22 week old male WT and MCT6-KO DIO mice following chronic administration of metformin (300 mg/kg, daily, p.o.). Additionally, KO + Lac-Phe group received daily administration of Lac-Phe (50 mg/kg, i.p.) while WT + Vehicle and KO + Vehicle groups received vehicle control (5% DMSO in saline). N = 8/group. Starting body weights were WT + Vehicle: 40.2 ± 1.8 g, KO + Vehicle: 41.0 ± 2.3 g, and KO + Lac-Phe: 40.8 ± 2.3 g (mean ± SEM). For (C-F) and (J-M), mice were on high fat diet for 8 weeks. For (G-I,N,O), mice were on high fat diet for 11-12 weeks. Data are shown as mean ± SEM. * *p* < 0.05, ** *p* < 0.01, *** *p* < 0.001. In (B,C,D,E,G,H,J), *p*-values were calculated with two-way ANOVA and reporting the effect of genotype. In (F), *p*-values were calculated with one-way ANOVA with post hoc Holm-Šídák’s multiple comparisons test. In (I), *p*-values were calculated with multiple unpaired t tests. In (K-M), *p*-values were calculated from two-tailed unpaired t-test. In (N,O), *p*-values were calculated from two-way ANOVA with post hoc Tukey’s multiple comparisons test.

Because of the established role for Lac-Phe in feeding and body weight regulation, we next assessed energy balance phenotypes in MCT6-KO mice. We first monitored the body weights and food intake of MCT6-KO and WT littermate controls on either chow or high fat diet (HFD, 60% kcal from fat). Under these conditions, no differences in body weights or food intake were observed (**Fig. S3F-I**). In metabolic chambers, another cohort of HFD-fed MCT6-KO mice exhibited identical ambulatory activity, VO2, VCO2, respiratory exchange ratio (RER), and food intake compared to WT controls (**Fig. S3J-N**). At the end of this experiment, tissues were harvested for molecular analysis. MCT6-KO mice did not exhibit any differences in mitochondrial or inflammation gene expression in liver, muscle or epididymal adipose tissues (**Fig. S3O-R**). MCT6-KO mice did not exhibit any large changes in a panel of plasma metabolic and lipid markers, including AST, ALT, and triglycerides (**Fig. S3S-U**), although we detected more minor reductions in HDL-c without changes to LDL-c or total cholesterol (**Fig. S3V-X**). Therefore, genetic ablation of MCT6 did not affect food intake or body weight following HFD feeding, nor did it grossly affect molecular measurements of metabolic function.

To introduce a glycolytic stimulus to increase Lac-Phe levels,^8,10,27^ we challenged a new cohort of HFD-fed, obese MCT6-KO mice with metformin by daily oral gavage (300 mg/kg/day, p.o.). Body weights were not different between genotypes prior to the metformin administration protocol (mean ± SEM: WT: 35.2 ± 1.1 g, MCT6-KO: 34.6 ± 1.5 g, *p* > 0.05). As expected,^8^ WT mice treated with metformin lost weight over the course of the experiment (mean ± SEM -0.8 ± 0.3 g). By contrast, MCT6-KO mice did not lose weight, and in fact gained a slight amount of weight over the same treatment period (mean ± SEM +0.3 ± 0.4 g, *p* < 0.01, **Fig. 4C**). Food intake was also higher in MCT6-KO mice compared to WT on metformin treatment (**Fig. 4D**). Both MCT6-KO and WT mice exhibited a normal glucose excursion in response to acute metformin treatment (**Fig. 4E**). Next, we pair-fed metformin-treated MCT6-KO mice to the food intake of metformin-treated WT mice (mean ± SEM: WT-metformin, 2.22 ± 0.06 g/day; KO-metformin, 2.42 ± 0.05 g/day; KO-metformin, pair-fed, 2.21 ± 0.07 g/day). As shown in **Fig. 4F**, pair-fed metformin-treated MCT6-KO mice lost weight to the same extent as metformin-treated WT animals, whereas metformin-treated MCT6-KO mice were once again resistant to drug-induced weight loss. We conclude that MCT6-KO mice exhibit resistance to the body weight-lowering effects of metformin treatment.

To further validate and extend these findings, we performed two additional experiments. First, we administered metformin (300 mg/kg/day, p.o.) to HFD-fed MCT6-KO and control mice for a longer period (1 month). Starting body weights were not different between genotypes (mean ± SEM: WT 37.1 ± 0.7 g; MCT6-KO 36.7 ± 1.4 g, *p* > 0.05). Again, we observed that MCT6-KO mice exhibited higher food intake and resistance to metformin-associated weight loss (**Fig. 4G,H**). Dissection of tissues at the end of this chronic 28-day experiment revealed greater adipose mass in MCT6-KO mice (**Fig. 4I**). The weights of the other organs were not different between genotypes (**Fig. 4I**). Second, in an independent cohort of HFD-fed MCT6-KO and control mice, we used a higher metformin dose (600 mg/kg/day, p.o.) to induce a more profound weight loss effect. Starting body weights were again not different between genotypes (mean ± SEM: WT 42.0 ± 1.4 g; MCT6-KO 40.0 ± 1.7 g, *p* > 0.05). We observed once again that WT mice lost more weight than MCT6-KO mice at this higher metformin dose (**Fig. 4J**). Blood levels of GDF15, GLP-1, and PYY were not different between WT and MCT6-KO mice after metformin administration (**Fig. 4K-M**). Therefore, the resistance of MCT6-KO mice to the body weight-lowering effects of metformin is not due to changes in these other feeding-associated hormones. The metformin effect on feeding and body weight was identical in MCT6-KO and WT mice on chow diet (**Fig. S3Y,Z**), establishing that the resistance to metformin-associated weight loss in MCT6-KO mice requires high fat diet feeding and/or obesity.

To directly test the causal role of Lac-Phe in the MCT6-KO mice, we performed a three-arm rescue experiment: WT mice treated with metformin, MCT6-KO mice treated with metformin, and MCT6-KO mice concurrently treated with metformin and Lac-Phe. As expected, WT mice exhibited reduced food intake and body weight on metformin treatment, whereas MCT6-KO mice were resistant (**Fig. 4N,O**). Concurrent Lac-Phe administration to MCT6-KO mice fully normalized the food intake and body weight back to that of metformin-treated WT mice (**Fig. 4N,O**). We conclude that bypassing the transporter defect with exogenous Lac-Phe is sufficient to normalize the body weight phenotype of metformin-treated MCT6-KO mice.

### Characterization of additional MCT6 substrates

To define the broader substrate scope of MCT6, we performed untargeted metabolomics on both media and cell lysates from MCT6-overexpressing (HEK293T and HCT116) and MCT6-KO (HCT116 KO1 and KO2) cells (see **Methods**). In total, we detected ∼2000 mass features in each media and cell lysate compartments from HEK293T cells, and ∼500 mass features in each compartment of HCT116 cells (**Table S2**). To isolate robust and reproducible MCT6-regulated metabolites, we retained only mass features that concordantly changed across replicate lines (elevated in both overexpression lines, or reduced in both knockout lines, in either media or cell lysate compartments). The full untargeted metabolomic datasets and mass features are provided in **Table S2**. In addition to Lac-Phe, our analysis identified the following additional mass features which were regulated by MCT6: *m/z* 252.09, retention time (RT) 7.5 min (M252T7); *m/z* 275.10, RT 6.7 min (M275T7); *m/z* 202.11, RT 5.8 min (M202T6); *m/z* 188.09, RT 6.7 min (M188T7); *m/z* 181.05, RT 6.9 min (M181T7); and *m/z* 190.05, RT 6.6 min (M190T7). By comparison of chromatographic elution and MS/MS spectra to authentic standards generated through independent chemical synthesis, we successfully determined their structures to be structurally related *N*-lactoyl amino acids Lac-Tyr (*N*-lactoyl-tyrosine), Lac-Trp (*N*-lactoyl-tryptophan), Lac-Leu/Ile (*N*-lactoyl-(iso)leucine), Lac-Val (*N*-lactoyl-valine), and, as well as two additional molecules, the lactate derivative *p*-hydroxyphenyllactic acid (HPLA) and *N*-acetylmethionine (Ac-Met) (**Fig. 5A** and **Fig. S4A-F**). Notably, many of these share a lactoyl chemical moiety (**Fig. 5A**). The levels of these metabolites in both media and cell lysate compartments are shown in **Fig. 5B** (overexpression comparisons) and **Fig. 5C** (knockout comparisons). In addition to Lac-Phe, three metabolites (Lac-Tyr, Lac-Trp, and Ac-Met) showed the most consistent and concordant reciprocal extracellular and intracellular changes across all four comparisons (e.g., higher in media and lower intracellularly in both HCT116 and HEK293T OE cells, and lower in media and higher intracellularly in HCT116 KO1 and KO2 cells). The other MCT6-regulated metabolites showed more changes that were restricted to a subset of conditions. For instance, Lac-Leu/Ile and Lac-Val were most changed in media of MCT6 overexpressing cells but not accumulated intracellularly in MCT6-KO cells (**Fig. 5B,C**). HPLA was most dramatically changed intracellularly but not extracellularly across all four comparisons (**Fig. 5B,C**). Using both *Xenopus* oocyte (**Fig. S4G-M**) and mammalian cell-based (**Fig. S4N-T**) transport assays, we confirmed all candidates were MCT6 substrates in vitro. Thus, in addition to Lac-Phe, MCT6 transports several other metabolites, including multiple additional *N*-lactoyl amino acids.

**Figure 5.**
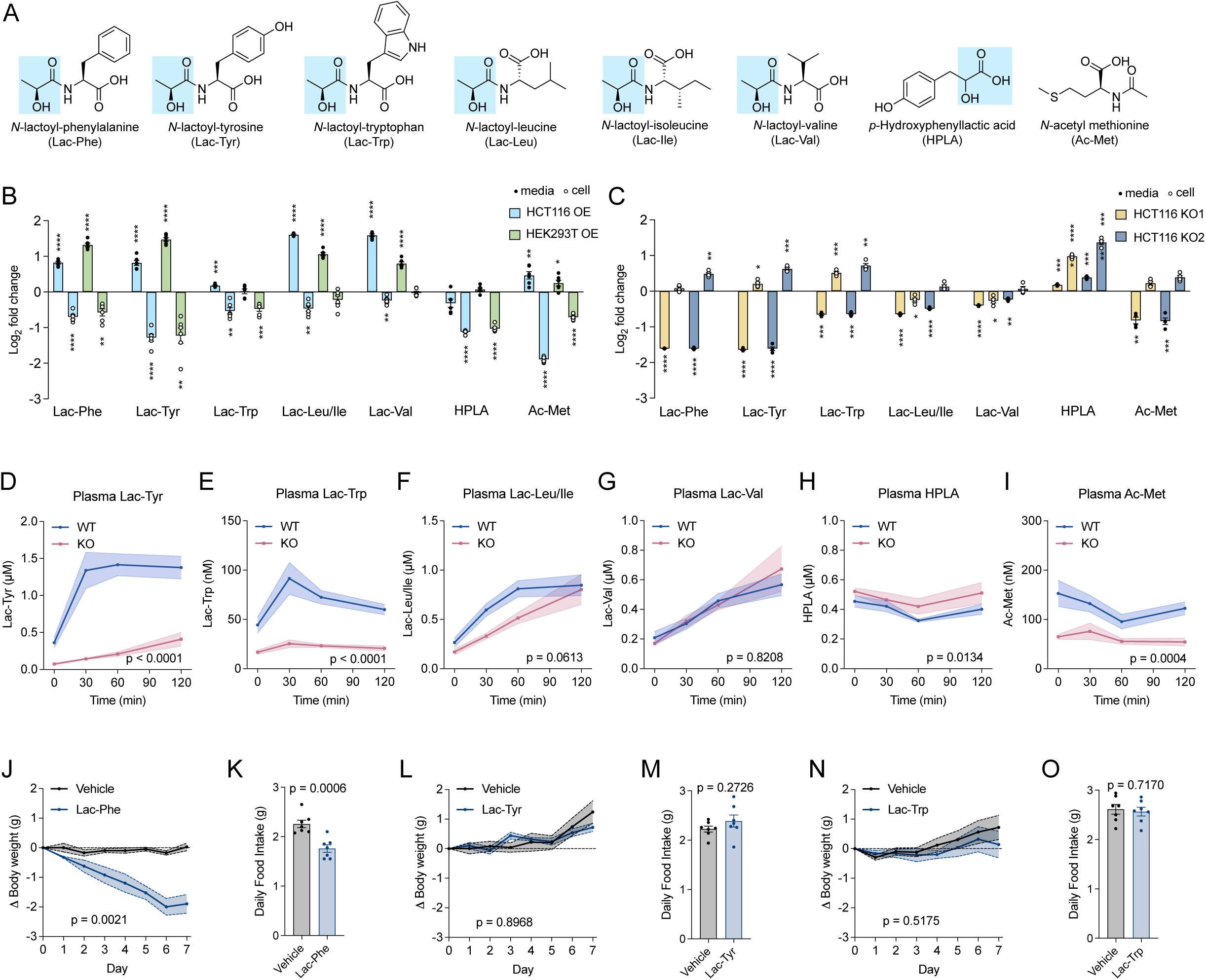
Characterization of additional MCT6 substrates. (A) Chemical structures of additional MCT6 substrates identified through untargeted profiling. (B) Levels of additional MCT6 substrates in conditioned media (filled circles) and cell lysates (open circles) in HCT116 or HEK293T cells overexpressing MCT6 (Dox+). Levels were normalized to control (Dox-) cells. N = 6/group. (C) Levels of additional MCT6 substrates in conditioned media (filled circles) and cell lysates (open circles) in HCT116 MCT6 KO cells. Levels were normalized to control cells. N = 4/group. (D-I) Plasma levels of additional MCT6 substrates at the indicated time point from 8-12 week old male WT and MCT6-KO mice before (time 0) and after a single administration of metformin (300mg/kg, p.o.). N = 7-8/group. (J,K) Cumulative change in body weight (J) and daily food intake (K) in 13-14 week old male C57BL6/J DIO mice following chronic administration of Lac-Phe (50 mg/kg daily, i.p.) or vehicle control. N = 4/group. (L,M) Cumulative change in body weight (L) and daily food intake (M) in 14-15 week old male C57BL6/J DIO mice following chronic administration of Lac-Tyr (50 mg/kg daily, i.p.) or vehicle control. N = 5/group. (N,O) Cumulative change in body weight (N) and daily food intake (O) in 14-15 week old male C57BL6/J DIO mice following chronic administration of Lac-Trp (50 mg/kg daily, i.p.) or vehicle control. N = 5/group. Data are shown as mean ± SEM. * *p* < 0.05, *** *p* < 0.001, **** *p* < 0.001. In (B,C), p-values were calculated from two-tailed one sample t tests. In (D-J,L,N), *p*-values were calculated from two-way ANOVA and reporting the effect of genotype or treatment. In (K,M,O), *p*-values were calculated with two-tailed unpaired t tests.

Next, we measured the levels of these additional metabolites from WT or global MCT6-KO mice at basal state and after metformin. In blood, only Lac-Tyr and Lac-Trp were both metformin-inducible and reduced in MCT6-KO mice (**Fig. 5D,E**). The other *N*-lactoyl amino acids, Lac-Leu/Ile and Lac-Val, also exhibited metformin inducibility, but were not significantly different between genotypes (**Fig. 5F,G**). Lastly, HPLA and Ac-Met were elevated and reduced, respectively, in MCT6-KO blood and did not exhibit regulation following metformin dosing (**Fig. 5H,I**). We also measured these same metabolites in the gut, muscle, liver, and brain in WT and MCT6-KO mice. In general, we did not observe any consistent differences across genotypes (**Fig. S5A-D**). We also profiled a broader metabolome panel across blood, as well as gut, muscle, liver, and brain tissues both in the basal state and after metformin treatment. No differences were observed between MCT6-KO and WT mice (**Fig. S5E-I**).

Two of these MCT6-regulated metabolites, Lac-Tyr and Lac-Trp, were metformin inducible and strongly reduced in MCT6-KO mice and might therefore plausibly contribute to the phenotype of MCT6-KO mice. However, their effects on feeding and body weight have not been previously evaluated. We synthesized large quantities of Lac-Tyr and Lac-Trp for in vivo administration to diet-induced obese mice (see **Methods**). Lac-Phe was used as a positive control. In this experiment, only Lac-Phe reduced body weight and feeding (**Fig. 5J,K**) whereas Lac-Tyr and Lac-Trp were both without effect (**Fig. 5L-O**). Thus, Lac-Tyr and Lac-Trp cannot explain the body weight phenotype of MCT6-KO mice.

### Effect of exercise in MCT6-KO mice

To determine whether MCT6 also regulates Lac-Phe during exercise, we first tested the effect of a moderate-intensity treadmill running protocol on the body weight and metabolite levels in MCT6-KO and WT mice on HFD (**Fig. S6A**, and see **Methods**). Under these conditions, we did not detect any differences between genotypes (**Fig. S6B,C**). To characterize the metabolite response under this same moderate-intensity exercise paradigm, we next measured circulating metabolites before and after a single bout of treadmill running. Basal circulating Lac-Phe was reduced by ∼50% in MCT6-KO mice (**Fig. S6D**), but following exercise, was increased to similar levels in both genotypes (**Fig. S6D**). The other MCT6-regulated metabolites were also induced by exercise (**Fig. S6E-J**). Lac-Tyr, Lac-Trp, and Ac-Met remained lower in MCT6-KO plasma, whereas Lac-Leu/Ile, Lac-Val, and HPLA did not differ between genotypes (**Fig. S6E-J**). None of these metabolites showed consistent genotype-dependent differences in gut, muscle, liver, or brain collected immediately after exercise (**Fig. S6K-N**), and broader metabolomic profiling similarly revealed no consistent tissue differences (**Fig. S6O-S**).

To determine whether the contribution of MCT6 to Lac-Phe levels is dependent on exercise intensity, we next used an extreme, maximal-intensity and high-incline sprint exercise challenge (**Fig. S6T**, and see **Methods**). Unlike in our prior exercise protocol (**Fig. S6A**), here MCT6-KO mice exhibited lower blood Lac-Phe levels both at baseline and all time points after the exercise bout (**Fig. S6U**). Lac-Tyr and Lac-Trp also remained lower in MCT6-KO plasma, whereas the other substrates were not different between genotypes (**Fig. S6V-AA**). Running time was also identical between genotypes (mean ± SEM: WT, 29.7 ± 1.6 min, MCT6-KO, 29.8 ± 1.7 min, *p* > 0.05). Maximal exertion of this kind cannot be sustained as a daily protocol and is therefore unsuitable for chronic body weight studies, for which metformin instead provided a tractable paradigm of sustained high Lac-Phe flux. *Slc16a5* mRNA levels trended higher in both gut and muscle after metformin or exercise, reaching statistical significance only for exercise in the gut (**Fig. S6AB,AC**). Thus, MCT6 is required for the exercise-inducible increases in circulating Lac-Phe during maximal, but not moderate-intensity, exercise.

### Intestine epithelium-specific deletion of MCT6 phenocopies the global MCT6-deficient mice

We originally identified MCT6 as a transporter co-expressed with CNDP2 in the gut. In the public expression database ProteomicsDB,^28^ mouse *Slc16a5* mRNA (encoding MCT6) is highly enriched in the gastrointestinal tract compared to other tissues (**Fig. S7A**). Furthermore, re-examination of intestine single cell RNAseq data confirmed co-expression of *Cndp2* and *Slc16a5* mRNA in multiple enterocyte populations across proximal and distal gut (**Fig. S7B**). To determine whether the phenotype of global MCT6-KO mice is due to intestinal epithelial expression of MCT6, we generated a conditional *Slc16a5* allele by flanking exon 1 with loxP sites (**Fig. 6A** and see **Methods**). We crossed this conditional allele to the intestinal epithelial Vil1-Cre driver to generate intestine epithelial-specific MCT6-KO mice (*Vil1^Cre+/-^; Slc16a5^fl/fl^*). Cre-negative, *Slc16a5^fl/fl^* mice were used as controls. *Vil1^Cre+/-^; Slc16a5^fl/fl^*animals exhibited selective reduction of *Slc16a5* mRNA in the intestine, with no changes in the kidney, liver, pancreas, or spleen (**Fig. 6B**). We also validated reduction of MCT6 protein in gut tissues from *Vil1^Cre+/-^; Slc16a5^fl/fl^* mice using an anti-MCT6 antibody (**Fig. 6C**). The residual band may potentially reflect gut MCT6 expression in non-epithelial cells (**Fig. 6C**). Gut CNDP2 protein levels were unchanged in *Vil1^Cre+/-^; Slc16a5^fl/fl^*mice (**Fig. 6C**). Next, we assessed the biochemical phenotype of *Vil1^Cre+/-^; Slc16a5^fl/fl^* mice. First, while baseline plasma Lac-Phe levels were not different between genotypes, metformin-stimulated plasma Lac-Phe was reduced by >50% at all post-metformin time points in *Vil1^Cre+/-^; Slc16a5^fl/fl^* mice (**Fig. 6D**). Some of the additional MCT6-regulated metabolites, especially the *N*-lactoyl amino acids, were also metformin inducible and reduced in *Vil1^Cre+/-^; Slc16a5^fl/fl^* mice (**Fig. S7C-H**). A broader metabolite profiling of blood revealed that most other metabolites remained unchanged between *Vil1^Cre+/-^; Slc16a5^fl/fl^* versus control mice (**Fig. 6E**).

**Figure 6.**
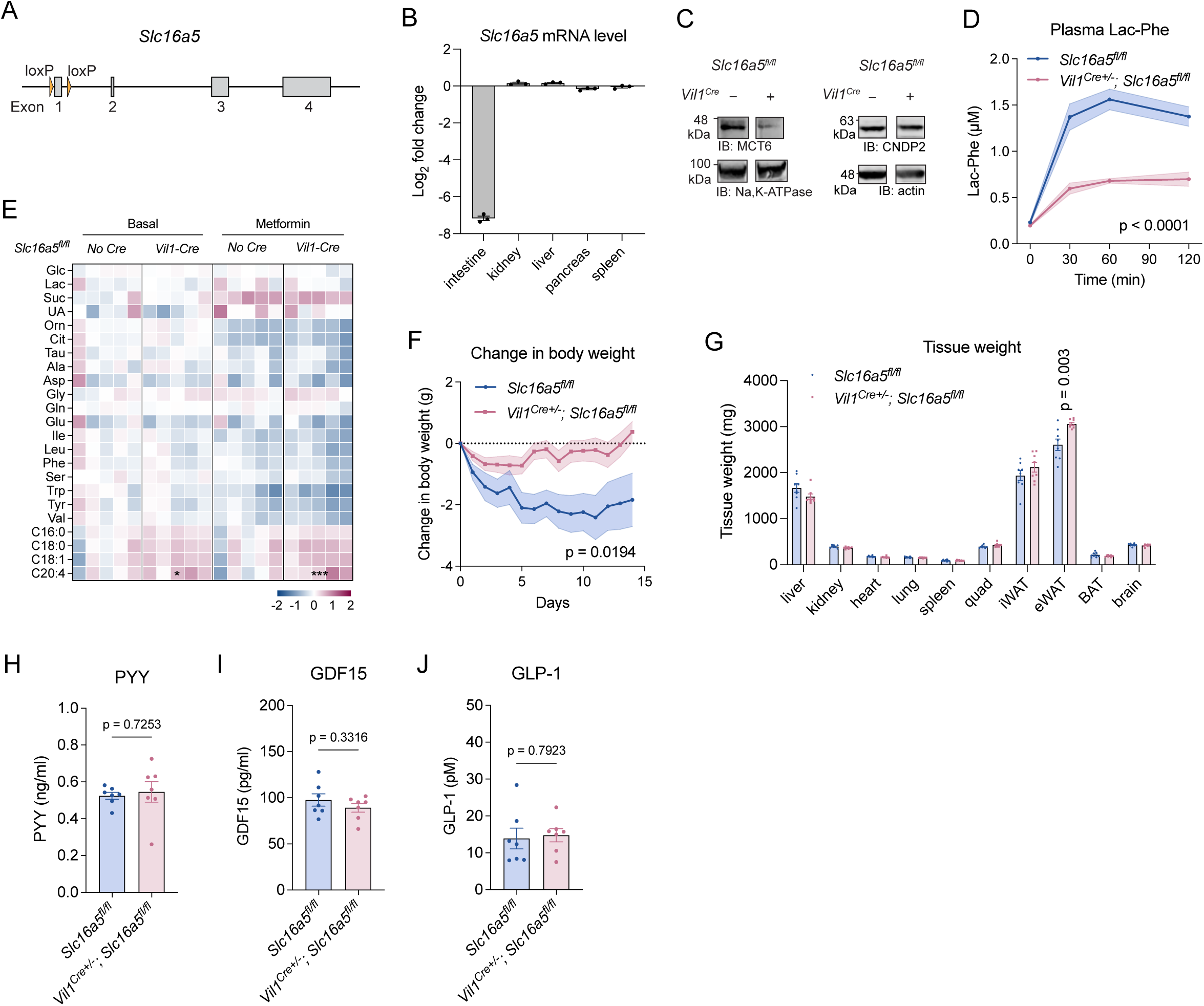
Characterization of intestinal epithelial-specific MCT6-KO mice. (A) Schematic illustration of the floxed allele of *Slc16a5* encoding MCT6. (B) Change of *Slc16a5* mRNA levels in tissues of *Vil1^Cre+/-^; Slc16a5^fl/fl^* mice compared to *Slc16a5^fl/fl^* controls. (C) Anti-MCT6 or anti-sodium potassium ATPase blotting (left) and anti-CNDP2 or anti-beta-actin blotting (right) of intestine lysates of *Vil1^Cre+/-^; Slc16a5^fl/fl^* mice and *Slc16a5^fl/fl^* controls. (D) Plasma Lac-Phe levels at the indicated time point from 7-8 week-old male *Vil1^Cre+/-^; Slc16a5^fl/fl^* mice and *Slc16a5^fl/fl^* controls before (time 0) and after metformin administration (300mg/kg, p.o.). N = 5-6/group. (E) Levels of metabolites in the plasma under basal state or 30 min after a single administration of metformin (300 mg/kg, p.o.). Levels were normalized to Cre-negative controls under basal state, with log_2_ transformation. Glc, glucose; Lac, lactate; Suc, succinate; UA, uric acid; Orn, ornithine; Cit, citrulline; Tau, taurine; Ala, alanine; Asp, aspartic acid; Gly, glycine; Gln, glutamine; Glu, glutamic acid; Ile, isoleucine; Leu, leucine; Phe, phenylalanine; Ser, serine; Trp, tryptophan; Tyr, tyrosine; Val, valine; C16:0, palmitate; C18:0, stearate; C18:1, oleate; C20:4, arachidonate. (F) Cumulative change in body weight in 14-18 week-old male *Vil1^Cre+/-^; Slc16a5^fl/fl^* DIO mice and *Slc16a5^fl/fl^* controls following chronic metformin treatment (300 mg/kg daily, p.o.). N = 8 per group. Starting body weights were *Vil1^Cre+/-^; Slc16a5^fl/fl^*: 40.4 ± 0.8 g, *Slc16a5^fl/fl^*: 42.8 ± 1.8 g (mean ± SEM). (G) Tissue weights of male *Vil1^Cre+/-^; Slc16a5^fl/fl^* DIO mice and *Slc16a5^fl/fl^*controls after chronic metformin treatment for 14 days. N = 8/group. (H-J) Plasma PYY (H), GDF15 (I), or GLP-1 (J) levels in 10-13 week old male *Vil1^Cre+/-^; Slc16a5^fl/fl^* mice and *Slc16a5^fl/fl^*controls 4 hours after a single administration of metformin (300 mg/kg, p.o.). N = 6-7/group. Data are shown as mean ± SEM. * *p* < 0.05, *** *p* < 0.001. In (D,F) *p*-values were calculated from two-way ANOVA and reporting the effect of genotype or treatment. In (E), *p*-values were calculated with two-way ANOVA with post hoc Šídák’s multiple comparisons test. In (G), *p*-values were calculated with multiple unpaired t tests. In (H-J), *p*-values were calculated from two-tailed unpaired t-test.

Lastly, we determined the body weight phenotypes of *Vil1^Cre+/-^; Slc16a5^fl/fl^* mice in response to chronic metformin treatment. *Vil1^Cre+/-^; Slc16a5^fl/fl^* mice were first rendered obese by high fat diet feeding for 8 weeks and then treated with metformin (300 mg/kg/day, p.o.) daily for two weeks. Prior to metformin treatment, body weight was not different between genotypes (mean ± SEM: *Vil1^Cre+/-^; Slc16a5^fl/fl^*: 40.4 ± 0.8 g, *Slc16a5^fl/fl^*: 42.8 ± 1.8 g, *p* > 0.05). However, upon treatment with metformin, *Vil1^Cre+/-^; Slc16a5^fl/fl^* mice exhibited resistance to metformin-induced weight loss (**Fig. 6F**) and had higher fat mass at the end of the two-week treatment, particularly in the epididymal white adipose (eWAT) depot, compared to control *Slc16a5^fl/fl^*mice (**Fig. 6G**). Once again, plasma levels of PYY, GDF15, and GLP-1 were not different between control and *Vil1^Cre+/-;^ Slc16a5^fl/fl^* mice after metformin administration (**Fig. 6H-J**). We conclude that intestinal epithelial MCT6 regulates metformin-inducible Lac-Phe levels and is required for metformin-associated weight loss.

## Discussion

Here we identify MCT6 as a physiologic intestinal Lac-Phe exporter and establish its role in metformin-associated weight loss. Multiple lines of evidence support this conclusion: 1) MCT6 is a gut-enriched transporter that exhibits high co-expression with the Lac-Phe biosynthetic enzyme CNDP2; 2) MCT6 is necessary and sufficient for Lac-Phe efflux in cell culture and directly transports Lac-Phe in vitro; 3) global and gut-specific MCT6-KO mice exhibit reduced metformin-inducible circulating Lac-Phe and resistance to metformin-associated weight loss; and 4) administration of exogenous Lac-Phe, which bypasses the export step, normalizes the body weight phenotype of MCT6-KO mice.

Blood-borne metabolite effectors present a distinctive cell biological problem in physiology. Unlike peptide hormones and neurotransmitters, many peripheral metabolites are not stored in secretory vesicles and must cross the plasma membrane to enter the circulation. Although transporter-dependent release has been described for several circulating effectors, rigorous genetic evidence that transporter ablation disrupts both circulating abundance and downstream systemic action remains rare.^3–5^ Our findings establish this causal sequence for Lac-Phe and underscore how Lac-Phe release to blood is not simply an automatic consequence of intracellular metabolite production, but in fact an independent and genetically encoded determinant. In addition, the biochemical segregation of extracellular versus intracellular pools raises the possibility that circulating metabolites like Lac-Phe have independent intracellular biological activity via engagement of intracellular targets. The lack of tissue accumulation of Lac-Phe in MCT6-KO mice might be explained by compensatory intracellular degradation pathways. We speculate that similar release-dependent control may extend to many additional blood-borne small molecule effectors.

Although MCT6 can transport a variety of xenobiotics in vitro,^18^ it has remained an “orphan” transporter lacking well-established physiologic substrates in vivo. We show that Lac-Phe is a bona fide physiologic MCT6 substrate. MCT6-mediated transport is Na⁺/Cl⁻-independent, pH-sensitive, and influenced by membrane electrochemical state, while mutagenesis identifies Gly347 and Arg288 as important for full transport activity. Lastly, for MCT6 we measure an apparent in vitro Lac-Phe *K*_m_ of ∼2.7 mM. This measured *K*_m_ is well above our bulk intracellular Lac-Phe levels (∼10-20 µM). Nevertheless, considering our genetic evidence of physiologic Lac-Phe transport by MCT6, one interpretation of this numerical mismatch is that our in vitro transport assays do not fully recapitulate the state of MCT6 in cells. Alternatively (or in addition), there may also be spatial interaction/coupling between CNDP2 and MCT6 which would raise the local substrate concentration of Lac-Phe at the transporter above bulk cellular average levels.

The identification of Lac-Phe as an MCT6 substrate also connects MCT6 with the pharmacology of metformin, a widely prescribed anti-diabetic medicine that also can reduce body weight in humans.^29^ In the Diabetes Prevention Program (DPP),^30,31^ average metformin-associated weight loss was -2.1 kg (baseline, 94.2 kg). Furthermore, this weight loss response is highly heterogeneous: 29% of metformin-treated participants lost ≥ 5% body weight at one year, and 8-10% lost ≥ 10% body weight within the first two years.^30^ The mechanisms underlying metformin-associated weight loss remain debated.^8,32,33^ Previously, we showed that genetic ablation of the Lac-Phe biosynthetic enzyme CNDP2 reduced circulating Lac-Phe and conferred resistance to metformin-associated weight loss.^8^ Here, disruption of Lac-Phe export to blood produces the same phenotype. Thus, two orthogonal genetic perturbations—one blocking Lac-Phe synthesis and the other limiting its entry into blood—support a necessary role for this pathway under our experimental conditions. The MCT6 phenotype is expressed as an altered change in body weight following metformin treatment rather than a difference in absolute body weight. Furthermore, the absolute magnitude of the metformin effect on body weight in our mouse studies was modest, consistent with the similarly modest (∼2-4%) effect of metformin on human body weight. Pair feeding shows that our genotype effect is explained by food intake, consistent with the anorexigenic activity of Lac-Phe^8^ and with prior studies implicating reduced energy intake in metformin-associated weight loss.^29^ An important role for the gut in metformin action, as illustrated by our gut-specific MCT6-KO mice, is also consistent with recent data from Sebo et al.^27^ The reproducibility of the phenotype across global and intestinal epithelial-specific knockout models, treatment durations, and metformin doses, together with rescue by exogenous Lac-Phe, further support this conclusion. Our findings do not, however, exclude contributions from other metformin-responsive pathways. For instance, the GDF15/GFRAL pathway has also been implicated in metformin-associated weight loss,^32,33^ although data in this area have been conflicting.^34^ Prior mediation analysis in the MESA cohort identified effects of both Lac-Phe and GDF15.^8^ These data, taken together, are consistent with a model in which metformin engages multiple anorexigenic signals whose relative contributions may vary across experimental settings and among individuals.

The contribution of MCT6 to circulating Lac-Phe also depends on stimulus intensity. MCT6 is required for the full Lac-Phe response to metformin and maximal sprint exercise but is dispensable for the increase during moderate treadmill exercise. One potential explanation is that MCT6 transport activity increases at acidic pH, so that export is stimulated by the same metabolic state that drives Lac-Phe synthesis. A conceptually related mechanism has been previously described for succinate.^35^ Under conditions of lower glycolytic flux or otherwise lower reductive stress, additional MCT6-independent routes are likely operational. Lastly, additional Lac-Phe transporters may also operate in other tissues.

MCT6 transports a broader set of endogenous metabolites, including several *N*-lactoyl amino acids. The enrichment of aromatic and hydrophobic *N*-lactoyl amino acids regulated by MCT6, particularly Lac-Tyr and Lac-Trp in vivo, is consistent with the predicted hydrophobicity of the MCT6 substrate-binding cavity. Among the metabolites altered in MCT6-KO blood, however, Lac-Phe uniquely combines metformin inducibility, MCT6 dependence, and anorexigenic activity. Lac-Tyr and Lac-Trp are induced by metformin and reduced in MCT6-KO mice but do not suppress feeding or body weight. Lastly, two prior ‘omics’ studies identified pathway-level associations with MCT6 loss but overt functional defects in metabolic homeostasis were not established.^19,20^ Notably, several of the most robustly reduced metabolites reported in Ren et al.^19^ were 1-carboxyethyl amino acids—subsequently reassigned as *N*-lactoyl amino acids^13^— independently corroborating the broader MCT6 substrate signature identified here, and which we now mechanistically attribute to MCT6-mediated transport.

There remain some limitations of this work and fertile areas for potential future research. First, reconstitution of MCT6-KO mice with a transport-deficient MCT6 mutant would provide the most direct in vivo test that the phenotype depends on transport activity itself. We are in the process of generating such knock-in animals to test this possibility. Second, additional biochemical studies, such as MCT6 reconstitution studies using proteoliposomes, or structural studies of MCT6 in complex with Lac-Phe as a ligand, would enable additional understanding and rationalization of the precise mechanism of transport. Third, the identity of the transporters that mediate Lac-Phe release particularly at lower glycolytic flux rates remains an important question for future work. Finally, the physiologic functions of the other MCT6-regulated *N*-lactoyl amino acid metabolites require study in contexts beyond energy balance.

## Methods

### Chemicals

The following chemicals were purchased from Millipore Sigma: dithiothreitol (D0632), doxycycline (D9891), 2,4-dinitrophenol (DNP, D198501), rotenone (R8875), bumetanide (B3023), p-hydroxyphenyllactic acid (H3253), potassium bicarbonate (237205), HEPES (H3375), MES (475893), calcium D-gluconate (G4625), sodium D-gluconate (S2054), N-acetyl methionine (01310), sodium L-lactate (L7022), ammonium acetate (A2706), Kolliphor EL (C5135), D-mannitol (M9647). The following chemicals were purchased from Cayman Chemical: metformin hydrochloride (13118), D-(+)-Glucose-^13^C_6_ (26707), valinomycin (10009152), nigericin (11437). The following chemicals were purchased from Thermo Fisher Scientific: HALT Protease inhibitor cocktail (78438), blasticidin (A1113903), oligomycin (AAK61898MA), *N*-methyl-D-glucamine (NMDG, AAL1428230), ammonium hydroxide (A669), L-phenylalanine (A13238), DMSO (BP231), ponceau S staining solution (A40000279). Magnesium D-gluconate (sc-221868) and potassium D-gluconate (sc-202565) were purchased from Santa Cruz Biotechnology. *N*-dodecyl-β-d-maltopyranoside (DDM, D310LA) was purchased from Anatrace. Polyethylenimine (PEI, 23966) was purchased from Polysciences.

### Synthesis of *N*-lactoyl amino acids

Lac-Phe was synthesized as described previously.^6^ LC-MS: 236.093 [M-H]^−^. ^1^H NMR (400 MHz, D_2_O) δ 7.33–7.19 (m, 5H), 4.65 (dd, *J* = 9.1, 5.2 Hz, 1H), 4.10 (q, *J* = 6.9 Hz, 1H), 3.23 (dd, *J* = 13.9, 5.2 Hz, 1H), 2.99 (dd, *J* = 13.9, 9.2 Hz, 1H), 1.10 (d, *J* = 7.0 Hz, 3H). Lac-Tyr was synthesized by Acme Biosciences Inc. with > 95% purity as follows: *L*-tyrosine methyl ester hydrochloride (1.0 eq.) was dissolved in dichloromethane (DCM, 0.256 M) and mixed with 1-ethyl-3-(3-dimethylaminopropyl)carbodiimide (EDCI, 1.5 eq.), 1-hydroxybenzotriazole (HOBt, 1.5 eq.), *L*-lactic acid (1.0 eq.) and *N*,*N*-diisopropylethylamine (DIEA, 3.0 eq.) at 0°C. The mixture was stirred at room temperature for 4 hours, diluted with DCM (100 mL), washed with water, brine and dried with sodium sulfate. After filtration and evaporation of the solvent, the crude product was purified by silica gel chromatography (100-200 silica gel, petroleum ether/ethyl acetate, 64/36) to give *N*-lactoyl tyrosine methyl ester. The above ester (1.0 eq.) was dissolved in tetrahydrofuran (THF)/H_2_O (14/7 mL, 0.25 M) and mixed with LiOH (2.0 eq.). The reaction mixture was stirred at room temperature for 1 hour and monitored by LC-MS. After completion, the mixture was acidified with 1 M HCl to pH 3. The solution was concentrated and purified with prep-HPLC (H_2_O with 0.1% trifluoroacetic acid (TFA) and acetonitrile, 58/42) to give Lac-Tyr. LC-MS: 252.088 [M-H]^−^. ^1^H NMR (400 MHz, DMSO-*d_6_*) *δ* 9.18 (s, 1H), 7.48 (d, *J* = 8.4 Hz, 1H), 6.93–6.91(m, 2H), 6.63–6.61 (m, 2H), 5.57 (d, *J* = 5.2 Hz, 1H), 4.41–4.36 (m, 1H), 3.93–3.89 (m, 1H), 2.95–2.85 (m, 2H), 1.11 (d, *J* = 6.8 Hz, 3H).

Lac-Trp was synthesized by Acme Biosciences Inc. with > 95% purity as follows: *L*-tryptophan benzyl ester (1.0 eq.) was dissolved in DCM (0.34 M) and mixed with *L*-lactic acid (3.0 eq.), EDCI (1.5 eq.), HOBt (1.5 eq.) and DIEA (3.0 eq.). The reaction mixture was stirred at room temperature overnight under N_2_. The reaction mixture was diluted with 300 mL water and extracted with 3 x 200 mL ethyl acetate. The combined organic layers were washed with brine, dried over anhydrous sodium sulfate. After filtration, the filtrate was concentrated under reduced pressure. The residue was purified by silica gel column chromatography eluting with (petroleum ether/ethyl acetate, 1/0 to 3/1) to afford *N*-lactoyl tryptophan benzyl ester. The above ester (1.0 eq.) was dissolved in methanol (0.27 M) and mixed with 10% Pd/C (10% w/w) and triethylamine (2.0 eq.). The reaction mixture was stirred at room temperature for 12 hours under H_2_ (15 psi). The reaction mixture was filtrated and concentrated, triturated with DCM and dried to afford Lac-Trp. LC-MS: Lac-Trp, 275.104 [M-H]^−^. ^1^H NMR (400 MHz, DMSO-*d_6_*) *δ* 10.89 (s, 1H), 8.30 (s, 1H), 7.58 (d, *J* = 7.6 Hz, 1H), 7.50 (d, *J* = 8.0 Hz, 1H), 7.31 (d, *J* = 8.0 Hz, 1H), 7.10 (s, 1H), 7.03 (t*, J* = 7.6 Hz, 1H), 6.93 (t, *J* = 7.6 Hz, 1H), 4.42 (q, *J* = 6.4 Hz, 1H), 3.94 (q, *J* = 6.8 Hz, 1H), 3.26 (dd, *J* = 14.4, 5.2 Hz, 1H), 3.20–3.11 (m, 2H), 1.15 (d, *J* = 6.8 Hz, 3H).

Lac-Leu, Lac-Ile, and Lac-Val were synthesized as follows: sodium *L*-lactate (1.2 eq.) was dissolved in dichloromethane (0.2 M) and treated with 3-[bis(dimethylamino)methyliumyl]−3*H*-benzotriazol-1-oxide hexafluorophosphate (HBTU, 1.2 eq.) at 0°C under argon. After 15 min, leucine methyl ester hydrochloride, isoleucine methyl ester hydrochloride, or valine methyl ester hydrochloride (1.0 eq. depending on the Lac-AA being synthesized) and DIEA (3.0 eq.) in DCM (0.2 M) was added to the mixture. The reaction was stirred for 16 hours under argon at ambient temperature. One-third of the solvent was removed and the DCM solution was washed with 5% HCl, 5% NaHCO_3_ and saturated NaCl solutions. The organic layer was dried (MgSO_4_), filtered and concentrated. The resulting crude product was purified by column chromatography, eluting with ethyl acetate/hexane to afford the *N*-lactoyl amino acid methyl esters. The above ester (1.0 eq.) was dissolved in THF (0.5 M) and treated with LiOH (2.0 eq.) in water (0.5 M). The solution was stirred at ambient temperature for 2 hours, and the solvent removed. The resulting residue was dissolved in DCM and acidified by 5% HCl to pH 3. The resulting mixture was extracted with ethyl acetate three times, and the combined organic layers were washed with saturated NaCl solution. The organic layer was dried (MgSO_4_), filtered, and concentrated. The resulting crude product was purified to recrystallization with ethyl acetate/hexane to give the *N*-Lactoyl amino acid products. LC-MS: Lac-Leu, 202.109 [M-H]^−^; Lac-Ile, 202.109 [M-H]^−^; Lac-Val, 188.093 [M-H]^−^. ^1^H NMR: Lac-Leu, (400 MHz, D_2_O) δ 4.45 (dd, *J* = 9.6, 4.7 Hz, 1H), 4.36–4.28 (m, 1H), 1.81–1.63 (m, 3H), 1.39 (d, *J* = 6.9 Hz, 3H), 0.93 (dd, *J* = 16.3, 6.1 Hz, 6H); Lac-Ile, (400 MHz, CDCl_3_) δ 4.33 (dd, *J* = 6.3, 2.9 Hz, 2H), 1.99 (d, *J* = 6.7 Hz, 1H), 1.55–1.43 (m, 1H), 1.39 (d, *J* = 6.9 Hz, 3H), 1.32–1.19 (m, 1H), 0.95 (d, *J* = 23.7 Hz, 6H); Lac-Val, (400 MHz, CDCl_3_) δ 4.39–4.26 (m, 2H), 2.31–2.19 (m, 1H), 1.40 (d, *J* = 7.0 Hz, 3H), 0.98 (t, *J* = 6.6 Hz, 6H).

Deuterated *N*-lactoyl amino acids (D_3_-Lac-AAs) were synthesized by Acme Biosciences Inc. with > 95% purity as follows: ethyl 2-(benzyloxy)acetate (1.0 eq.) was dissolved in THF (0.645 M) at -78°C and mixed with 1 M lithium bis(trimethylsilyl)amide (LiHMDS, 1.0 eq.). The reaction mixture was stirred at -78°C for 30 min. Then CD_3_I (1.0 eq.) was added at -78°C. The reaction mixture was stirred at 0°C for 1 h. The reaction was monitored by TLC and LC-MS. After completion, the reaction was poured into aq. NH_4_Cl and extracted with ethyl acetate three times. The combined organic layer was washed with brine, dried over Na_2_SO_4_ and concentrated under reduced pressure. The residue was purified by column chromatography (100-200 silica gel, 20% ethyl acetate in petroleum ether as eluent) to give the desired product ethyl 2-(benzyloxy)propanoate-3,3,3-d_3_. The above product (1.0 eq.) was dissolved in MeOH/H_2_O (600 mL/200 mL, 0.384 M) and mixed with NaOH (1.5 eq.) at 0°C. The reaction mixture was stirred at room temperature for 2 hours. The reaction was monitored by TLC and LC-MS. After completion, the reaction mixture was acidified by 1 M HCl to pH 4, poured into water, and extracted with ethyl acetate three times. The combined organic layer was washed with brine, dried over Na_2_SO_4_ and concentrated under reduced pressure to give the desired product 2-(benzyloxy)propanoic-3,3,3-d_3_ acid. The above product (1.0 eq.) was dissolved in DCM (0.612 M) and mixed with dimethylformamide (DMF, 0.01 eq.) and oxalyl chloride (1.5 eq.) at 0°C. The reaction mixture was stirred at room temperature for 2 hours. The reaction was monitored by TLC and LC-MS. After the reaction was completed, the reaction was concentrated under reduced pressure to give the desired product 2-(benzyloxy)propanoyl-3,3,3-d_3_ chloride. (*R*)-4-benzyl-2-oxazolidinone (1.0 eq.) was dissolved in THF (0.297M) and mixed with *n*-BuLi (2.5 M in THF, 1.0 eq.) at -78°C. The reaction mixture was stirred at -78°C for 30 min. Then the above product (1.0 eq.) was added at -78°C. The reaction mixture was stirred at 0°C for 1 hour. The reaction was monitored by TLC and LC-MS. After the reaction was completed, the reaction was poured into aq. NH_4_Cl and extracted with ethyl acetate for three times. The combined organic layer was washed with brine, dried over Na_2_SO_4_ and concentrated under reduced pressure. The residue was purified by column chromatography (100-200 silica gel, 40% ethyl acetate in petroleum ether as eluent) to give the desired diastereomer (*R*)-4-benzyl-3-((*S*)-2-(benzyloxy)propanoyl-3,3,3-*d*_3_)oxazolidin-2-one. The product (1.0 eq.) was dissolved in THF/H_2_O (300/150 mL, 0.298 M) and mixed with 30% H_2_O_2_ (3.0 eq.) and LiOH·H_2_O (2.0 eq.) at 0°C. The mixture was stirred at room temperature for 4 hours. The reaction was monitored by TLC and LC-MS. After the reaction was completed, the reaction was poured into water and extracted with ethyl acetate three times. The combined organic layer was washed with brine, dried over Na_2_SO_4_ and concentrated under reduced pressure. The residue was purified by column chromatography (100-200 silica gel, 8% MeOH in DCM as eluent) to give the desired product (*S*)-2-(benzyloxy)propanoic-3,3,3-*d*_3_ acid. The product was dissolved in DCM (0.546 M) and mixed with *N*-hydroxysuccinimide (HOSU, 1.5 eq.) and EDCI·HCl (1.5 eq.) at room temperature. The reaction mixture was stirred at room temperature for 2 hours under N_2_. The reaction was monitored by TLC and LC-MS. After the reaction was completed, the reaction was poured into water and extracted with DCM three times. The combined organic layer was washed with brine, dried over Na_2_SO_4_ and concentrated under reduced pressure. The residue was purified by column chromatography (100-200 silica gel, 25% ethyl acetate in petroleum ether as eluent) to give the desired product *d*_3_-Lac(OBn)-ONSu.

For D_3_-Lac-Phe synthesis, *d*_3_-Lac(OBn)-ONSu (1.0 eq.) was dissolved in DMF (1 M) and mixed with DIEA (1.5 eq.) and *L*-phenylalanine (1.0 eq.) at 0°C. The reaction mixture was stirred at room temperature for 1 hour. The reaction was monitored by TLC and LC-MS. After the reaction was completed, the reaction was poured into water and extracted with ethyl acetate three times. The combined organic layer was washed with brine, dried over Na_2_SO_4_ and concentrated under reduced pressure. The residue was purified by column chromatography (100-200 silica gel, 8% MeOH in DCM as eluent) to afford D_3_-Lac-Phe benzyl ester. The ester (1.0 eq.) was dissolved in ethyl acetate (0.484 M), and mixed with Pd/C (100% w/w) under H_2_. The mixture was stirred at room temperature for 6 hours under H_2_. The reaction was monitored by TLC and LC-MS. After the reaction was completed, the reaction was filtered and washed with DCM/MeOH (10/1). The filtrate was concentrated and dried under reduced pressure to afford D_3_-Lac-Phe. LC-MS: 239.112 [M-H]^−^. ^1^H NMR (400 MHz, DMSO-*d_6_*) δ7.57–7.55 (m, 1H), 7.25–7.12 (m, 6H), 5.52 (s, 1H), 4.47–4.42 (m, 1H), 3.87 (s, 1H), 3.13–2.94 (m, 2H).

For D_3_-Lac-Tyr synthesis, *d*_3_-Lac(OBn)-ONSu (1.0 eq.) was dissolved in DMF (0.5 M) and mixed with DIEA (1.5 eq.) and *L*-tyrosine (1.0 eq.) at 0°C. The reaction mixture was stirred at room temperature for 1 hour. The reaction was monitored by TLC and LC-MS. After the reaction was completed, the reaction was poured into water and extracted with ethyl acetate three times. The combined organic layer was washed with brine, dried over Na_2_SO_4_ and concentrated under reduced pressure. The residue was purified by column chromatography (100-200 silica gel, 10% MeOH in DCM as eluent) to afford D_3_-Lac-Tyr benzyl ester. The ester (1.0 eq.) was dissolved in ethyl acetate (0.653 M), and mixed with Pd/C (100% w/w) under H_2_. The mixture was stirred at room temperature for 6 hours under H_2_. The reaction was monitored by TLC and LC-MS. After the reaction was completed, the reaction was filtered and washed with DCM/MeOH (10/1). The filtrate was concentrated and dried under reduced pressure to afford D_3_-Lac-Tyr. LC-MS: 255.107 [M-H]^−^. ^1^H NMR (400 MHz, DMSO-*d_6_*) δ 9.16 (br, 1H), 7.46–7.44 (m, 1H), 6.92–6.86 (m, 2H), 6.62–6.56 (m, 2H), 5.53 (br, 1H), 4.37–4.32 (m, 1H), 3.88 (s, 1H), 2.96–2.82 (m, 2H).

For D_3_-Lac-Trp synthesis, *d*_3_-Lac(OBn)-ONSu (1.0 eq.) was dissolved in DMF (0.393 M) and mixed with DIEA (1.5 eq.) and *L*-tryptophan (1.0 eq.) at 0°C. The reaction mixture was stirred at room temperature for 1 hour. The reaction was monitored by TLC and LC-MS. After the reaction was completed, the reaction was poured into water and extracted with ethyl acetate three times. The combined organic layer was washed with brine, dried over Na_2_SO_4_ and concentrated under reduced pressure. The residue was purified by column chromatography (100-200 silica gel, 8% MeOH in DCM as eluent) to afford D_3_-Lac-Trp benzyl ester. The ester (1.0 eq.) was dissolved in ethyl acetate (0.523 M), and mixed with Pd/C (100% w/w) under H_2_. The mixture was stirred at room temperature for 6 hours under H_2_. The reaction was monitored by TLC and LC-MS. After the reaction was completed, the reaction was filtered and washed with DCM/MeOH (10/1). The filtrate was concentrated and dried under reduced pressure to afford D_3_-Lac-Tyr. LC-MS: 278.123 [M-H]^−^. ^1^H NMR (400 MHz, DMSO-*d_6_*) δ 12.8 (br, 1H), 10.84 (s, 1H), 7.55–7.46 (m, 2H), 7.30–7.28 (m, 1H), 7.06–6.93 (m, 3H), 5.52 (s, 1H), 4.52–4.47 (m, 1H), 3.91–3.90 (m, 1H), 3.16–3.14 (m, 2H).

For D_3_-Lac-Leu synthesis, *d*_3_-Lac(OBn)-ONSu (1.0 eq.) was dissolved in DMF (0.446 M) and mixed with DIEA (1.5 eq.) and *L*-leucine (1.0 eq.) at 0°C. The reaction mixture was stirred at room temperature for 1 hour. The reaction was monitored by TLC and LC-MS. After the reaction was completed, the reaction was poured into water and extracted with ethyl acetate three times. The combined organic layer was washed with brine, dried over Na_2_SO_4_ and concentrated under reduced pressure. The residue was purified by column chromatography (100-200 silica gel, 10% MeOH in DCM as eluent) to afford D_3_-Lac-Leu benzyl ester. The ester (1.0 eq.) was dissolved in ethyl acetate (0.539 M), and mixed with Pd/C (100% w/w) under H_2_. The mixture was stirred at room temperature for 6 hours under H_2_. The reaction was monitored by TLC and LC-MS. After the reaction was completed, the reaction was filtered and washed with DCM/MeOH (10/1). The filtrate was concentrated and dried under reduced pressure to afford D_3_-Lac-Leu. LC-MS: 205.127 [M-H]^−^. ^1^H NMR (400 MHz, DMSO-*d_6_*) δ7.61–7.59 (m, 1H), 4.23 (m, 1H), 3.93 (s, 1H), 1.57–1.49 (m, 3H), 0.86–0.81 (m, 6H).

For D_3_-Lac-Ile synthesis, *d*_3_-Lac(OBn)-ONSu (1.0 eq.) was dissolved in DMF (0.515 M) and mixed with DIEA (1.5 eq.) and *L*-isoleucine (1.0 eq.) at 0°C. The reaction mixture was stirred at room temperature for 1 hour. The reaction was monitored by TLC and LC-MS. After the reaction was completed, the reaction was poured into water and extracted with ethyl acetate three times. The combined organic layer was washed with brine, dried over Na_2_SO_4_ and concentrated under reduced pressure. The residue was purified by column chromatography (100-200 silica gel, 10% MeOH in DCM as eluent) to afford D_3_-Lac-Ile benzyl ester. The ester (1.0 eq.) was dissolved in ethyl acetate (0.561 M), and mixed with Pd/C (100% w/w) under H_2_. The mixture was stirred at room temperature for 6 hours under H_2_. The reaction was monitored by TLC and LC-MS. After the reaction was completed, the reaction was filtered and washed with DCM/MeOH (10/1). The filtrate was concentrated and dried under reduced pressure to afford D_3_-Lac-Ile. LC-MS: 205.127 [M-H]^−^. ^1^H NMR (400 MHz, DMSO-*d_6_*) δ7.45–7.42 (m, 1H), 5.62 (br, 1H), 4.21–4.17 (m, 1H), 3.96 (s, 1H), 1.81–1.78 (m, 1H), 1.40–1.35 (m, 1H), 1.16–1.10 (m, 1H), 0.86–0.84 (m, 6H).

For D_3_-Lac-Val synthesis, *d*_3_-Lac(OBn)-ONSu (1.0 eq.) was dissolved in DMF (0.515 M) and mixed with DIEA (1.5 eq.) and *L*-valine (1.0 eq.) at 0°C. The reaction mixture was stirred at room temperature for 1 hour. The reaction was monitored by TLC and LC-MS. After the reaction was completed, the reaction was poured into water and extracted with ethyl acetate three times. The combined organic layer was washed with brine, dried over Na_2_SO_4_ and concentrated under reduced pressure. The residue was purified by column chromatography (100-200 silica gel, 10% MeOH in DCM as eluent) to afford D_3_-Lac-Val benzyl ester. The ester (1.0 eq.) was dissolved in ethyl acetate (0.589 M), and mixed with Pd/C (100% w/w) under H_2_. The mixture was stirred at room temperature for 6 hours under H_2_. The reaction was monitored by TLC and LC-MS. After the reaction was completed, the reaction was filtered and washed with DCM/MeOH (10/1). The filtrate was concentrated and dried under reduced pressure to afford D_3_-Lac-Val. LC-MS: 191.112 [M-H]^−^. ^1^H NMR (400 MHz, DMSO-*d_6_*) δ7.41–7.39 (m, 1H), 6.60 (br, 1H), 4.16–4.12 (m, 1H), 3.96 (s, 1H), 2.09–2.01 (m, 1H), 0.88–0.79 (m, 6H).

### Antibodies

Mouse anti-FLAG (Sigma, F1804); rabbit anti-HA (Cell Signaling Technology, 3724); rabbit anti-MCT6 (Proteintech, 12120-1-AP, used in all cell line blots); rabbit anti-MCT6 (Invitrogen, PA5-75154, used in tissue lysate blots, 1:500); mouse anti-β-tubulin (Cell Signaling Technology, 86298S); mouse anti-α-tubulin (Cell Signaling Technology, 2144S); rabbit anti-β-tubulin (OriGene, TA3101569); mouse anti-β-actin (Sigma, A1978); mouse anti-sodium, potassium-ATPase antibody (Abcam, ab7671-50); goat anti-mouse IRDye 680RD (LICORbio, 926-68070); goat anti-rabbit IRDye 800CW (LICORbio, 926-32211); Alexa Fluor 546-conjugated goat anti-rabbit (Invitrogen, A11035). Primary antibodies were used at 1:1000 unless specified.

Secondary antibodies were used at 1:5000, unless specified.

### Cell culture

HCT116 and HEK293T cells were obtained from the American Type Culture Collection (ATCC) and grown in DMEM (Corning, 10-017-CV) with 10% fetal bovine serum (FBS, Corning 35-010-CV) and 1% Penicillin-Streptomycin (Pen-Strep, Gibco 15140) in a humidified incubator at 37 °C with 5% CO_2_.

### Mouse information

All animal experiments were performed in accordance with procedures approved by the Stanford University Administrative Panel on Laboratory Animal Care. Mice were maintained in 12-hour light-dark cycles at 22°C with 50% relative humidity and fed a standard irradiated rodent chow diet. Where indicated, a high-fat diet (60% kcal from fat, Research Diets, D12492) was used. C57BL/6J (000664) and C57BL/6J DIO (380050) mice were purchased from the Jackson Laboratory. MCT6-KO mice were obtained from Dr. Jason Sprowl, University at Buffalo. MCT6-KO mice and WT littermates used in this study were generated by breeding heterozygous mice. Progeny pups were genotyped with the following primers: forward, 5′- ATCTCTTAAGCCCCCGGCTA-3′; reverse, 5′-ATAAGCAGTTCCACCCACCC-3′. *Slc16a5* flox mice were generated by the Transgenic, Knockout, and Tumor Model Center at Stanford University. Exon 1 of *Slc16a5* was flanked by two loxP sites, introduced by CRISPR/Cas9-mediated homology-directed repair using the following gRNA sequences: 5′- CAGCCCTAAGTATCAAGGAG-3′ and 5′-GCTACACTGAGTCCTGTCGG-3′. The following genotyping primers were used: forward, 5′-GGAAACGCCAAGGCCACTGTG-3′; reverse, 5′- CCGTCTCCCATCCCTAATGC-3′. Vil1-Cre mice were purchased from the Jackson Laboratory(021504) and genotyped using the following primers: forward, 5′- GCCTTCTCCTCTAGGCTCGT-3′; reverse, 5′-AGGCAAATTTTGGTGTACGG-3′.

### Transfection and Lac-Phe transporter screening

HEK293T cells were plated into 12-wells at 0.2 million cells per well. Next morning, cells in each well were transfected with 1 µg plasmid DNA using 6 µL PEI, with fresh media replaced 6 hours later. Conditioned media and cell lysates were collected 24 hours after the media change for metabolite measurements by LC-MS. One well was used for Western blot to analyze the expression of the FLAG-tagged transporter.

### Generation of doxycycline-inducible MCT6 overexpression cell lines

We subcloned hMCT6 from pDONR221_SLC16A5 (Addgene #131923) into pCW57.1_TStrp_HA_Ct (Addgene #194066), a doxycycline-inducible lentiviral vector containing C-terminal HA-Twin-Strep tags [PMID: 32265506]. Both plasmids were a gift from RESOLUTE Consortium & Giulio Superti-Furga. Lentiviral particles were produced in HEK293T cells transfected using PEI with the cloned lentiviral vector plasmid, together with the viral packaging psPAX2 plasmid and the viral envelope pMD2.G plasmid. The conditioned medium was collected 48 hours later, filtered through a 0.45-µm filter, and mixed 1:1 with 16 µg/mL polybrene (final concentration 8 µg/mL). One day after lentiviral transduction, HEK293T and HCT116 cells were subjected to blasticidin selection (10 µg/ml) for 10 days. Transgene expression was induced by 24 hour doxycycline treatment at 1 μg/ml and validated by Western blot.

To measure metabolite levels, cells were plated at 0.5 million cells per well in 12-well plates. One day later, medium was replaced with fresh medium containing 1 µg/ml doxycycline (Dox +), or vehicle control (0.01% ethanol, Dox -). Conditioned media and cell lysates were collected one day later for metabolite analysis by LC-MS.

### Generation of MCT6-KO HCT116 cell lines

MCT6-KO HCT116 cells were generated using the pLentiCRISPRv2 system, a gift from Feng Zhang (Addgene plasmid, 52961) [PMID: 25075903]. The sequences of guide RNAs used were as follows: KO1, 5’- gcttctactttgtccgccgg-3’ and 5’- gttgtgccgcaggatgtcga-3’; KO2, 5’- tcctggtgccatatgccatg-3’ and 5’-gtacttgcggtggctagcaa-3’. Oligonucleotides for the sgRNA and reverse complement sequences were synthesized and cloned into the plentiCRISPRv2 vector. A plentiCRISPRv2 plasmid without any gRNA insert was used as the control. HCT116 cells were transduced with lentiviral particles as described above and subjected to puromycin selection (1 µg/mL) for 10 days. Knockout efficiency of MCT6 was determined by genomic DNA sequencing. Genomic DNA was isolated with QIAamp DNA Mini Kit (Qiagen, 51304), the sequenced region was amplified with Phusion high-fidelity PCR master mix (New England Biolabs, M0531S) using the following primers: forward: 5’-CTGGGCATGTGCTTCAGCTTCC-3’; reverse: 5’- GTACGCCAGGCAGTAGCCCACG-3’. DNA bands were gel purified and sequenced by Genewiz, Azenta. The knockout efficiency was determined with Inference of CRISPR Edits (ICE) by EditCo Bio.

To measure metabolite levels, cells were plated at 1 million cells per well in 12-well plates. Conditioned media and cell lysates were collected one day later for metabolite analysis by LC-MS. For metformin treatment, cells were plated at 0.5 million cells per well in 12-well plates, and treated with 0 or 5 mM metformin next day. Conditioned media and cell lysates were collected 16 hours later for metabolite analysis by LC-MS.

### Lactate and N-lactoyl amino acid tracing using ^13^C_6_-glucose

^13^C glucose or ^12^C glucose of the same molar concentration (25 mM) was added to glucose-free DMEM (Gibco, cat. no. A1443001) supplemented with 10% FBS, 1% Pen-Strep and 4 mM L-glutamine. Cells plated in 12-well plates (at a density of 1 million cells per well) were washed with PBS twice before culturing in DMEM containing ^13^C glucose or ^12^C glucose. Conditioned media and cell lysates were collected 16 hours later for metabolite analysis by LC-MS.

### Cell culture sample preparation for LC-MS

For a 12-well plate with 1 mL media per well, 800 µL of conditioned media was acidified with 40 µL 1M HCl and extracted with 800 µL ethyl acetate. Samples were mixed by vortexing for 30 seconds. 400 µL of the upper ethyl acetate layer was dried under a stream of nitrogen. After a complete dry, samples were resuspended with 200 µL acetonitrile:water (90:10, v/v). For cell lysates, after washing the cells twice with cold saline, the cells were directly lysed with 200 µL acetonitrile:water (90:10, v/v). For both conditioned media and cell lysates, the acetonitrile:water mixture was centrifuged at 4°C for 15 min at 21,130 × g and the supernatant was carefully transferred to an LC-MS vial.

### Cellular respiration measurements

Cellular oxygen consumption rates (OCR) and extracellular acidification rates (ECAR) were determined using an Agilent Seahorse XF96 Analyzer. The assay was performed following the Agilent Seahorse XF Cell Mito Stress Test Kit manual, with slight modifications. In brief, cells were plated at 0.1 million cells per well in the XF96 cell culture plate and incubated with or without metformin in growth media overnight. The next morning, the cells were washed with PBS and incubated with Seahorse assay buffer (8.3 g/L DMEM (Sigma, D5030), 1.8 g/L NaCl, 1 mM pyruvate, 20 mM glucose, 1% Pen-Strep, pH 7.4). Final concentrations of compounds were used as follows: oligomycin, 10 μM; DNP, 100 μM; rotenone, 3 μM.

### CNDP2 enzymatic activity in cell lysates

Cells were washed twice with cold PBS and harvested by scraping. Cell pellets were collected by centrifugation (100 x g, 4°C, 5 min) and resuspended in 1 mL ice-cold PBS. Cells were lysed by sonication, and lysates were clarified by centrifugation at 21,130 × g for 10 min at 4°C. Supernatants were collected, and protein concentration was determined by BCA assay and normalized to 2 mg/mL. Protein lysate (50 µL, 100 µg total protein) was mixed with 50 µL of 40 mM lactate and phenylalanine (20 mM final concentration each). As a negative control, a parallel set of lysates was heat-denatured (95°C, 10 min) prior to substrate addition. Reactions were briefly vortexed, centrifuged, and incubated at 37°C for 1 h. Reactions were quenched with 5 µL of 1 M HCl, and metabolites were extracted with 400 µL ethyl acetate. The organic (top) layer (300 µL) was dried under nitrogen and reconstituted in 200 µL of acetonitrile:water (90:10, v/v) for LC-MS analysis.

### Western blot analysis

Cultured cells were lysed with a lysis buffer containing 25 mM Tris-HCl pH 8.0, 150 mM NaCl, 1% *n*-dodecyl-β-d-maltopyranoside (DDM), and 1x HALT. Lysates were incubated with constant rotation for 1 hour at 4°C, and centrifuged at 21,130 x g for 30 minutes at 4°C. Protein concentration was normalized with BCA assay. Proteins were diluted with 4x NuPAGE LDS Sample Buffer (ThermoFisher, NP008) and DTT (final concentration 100 mM). Proteins were separated on NuPAGE 4-12% Bis-Tris gels and transferred to nitrocellulose membranes. Blots were incubated with Odyssey blocking buffer for 1 hour and incubated with primary antibodies overnight at 4°C. Blots were then washed 3 times with PBST (0.1% Tween-20 in PBS) and stained with species-matched secondary antibodies at room temperature for 1 hour. Blots were washed 3 times with PBST and imaged with the Odyssey Fc Imaging System.

### RNA isolation, cDNA synthesis, and qPCR

Total RNA was extracted from cells or tissues using TRIzol reagent following standard phenol-chloroform separation, and the aqueous phase was further purified using an RNeasy column (Qiagen) with on-column ethanol washes, per manufacturer’s instructions. cDNA was synthesized from 1 µg of total RNA using the High-Capacity cDNA Reverse Transcription Kit (Applied Biosystems/Fisher, 4368814) according to the manufacturer’s protocol. qPCR reactions were assembled in 384-well plates at a final volume of 8 µL per reaction, comprising 4 µL of primer mix (1 µM each of forward and reverse primer) and 4 µL of SYBR Green master mix combined with diluted cDNA at a 20:1 ratio. For quantification of human *SLC16A5* (MCT6) mRNA in HEK293T and HCT116 cells, with *ACTB* and *GAPDH* as reference genes, the following primers were used: *SLC16A5* pair 1, forward 5′-TCAGCCAGCTCTACTTCACAG-3′ and reverse 5′-CGCCGAAGACAAGGAAGGT-3′; pair 2, forward 5′- CTTGTCTTCGGCGGGATCTTT-3′ and reverse 5′-GGACCACATCACACCCAGTA-3′; pair 3, forward 5′-TGGACGCCACCAACAACTTTA-3′ and reverse 5′-CTGCCACCCATGAAGAGGG-3′. *ACTB*, forward 5′-CACCATTGGCAATGAGCGGTTC-3′ and reverse 5′- AGGTCTTTGCGGATGTCCACGT-3′. *GAPDH*, forward 5′-GTCTCCTCTGACTTCAACAGCG-3′ and reverse 5′-ACCACCCTGTTGCTGTAGCCAA-3′. For quantification of mouse *Slc16a5* mRNA in mouse tissues, with *Actb* and *Gapdh* as reference genes: *Slc16a5* pair 1, forward 5′- CACCTGCATCGGTGTCTTCTT-3′ and reverse 5′-AAGGAAACCACGAGGTCTCAC-3′; pair 2, forward 5′-CATCTTGGTCAAACATTTCGGC-3′ and reverse 5′-AGAAGGTGCTGACTACCATGC-3′. *Actb*, forward 5′-GGCTGTATTCCCCTCCATCG-3′ and reverse 5′- CCAGTTGGTAACAATGCCATGT-3′. *Gapdh*, forward 5′-AGGTCGGTGTGAACGGATTTG-3′ and reverse 5′-TGTAGACCATGTAGTTGAGGTCA-3′.

#### Transporter assay in *Xenopus laevis* oocytes

Mouse *Slc16a5* cDNA sequence was codon optimized for expression in *Xenopus* and cloned into backbone pGEMHE [PMID: 1419000]. The plasmid was linearized with NheI for in vitro transcription with the mMESSAGE mMACHINE kit (Thermo Scientific, AM1344). The size and integrity of cRNAs were validated using gel electrophoresis (Reliant RNA Gels; Lonza, 54922). Each oocyte received an injection of 30 ng cRNA. Water-injected oocytes were used as controls. Injected *Xenopus* oocytes were incubated in 24-well plates at 17-18°C in modified L-15 oocyte media for 3-4 days before transporter assays. The L-15 oocyte media contains (per 1 L) 7.4 g Leibovitz’s Medium L-15 powder (Sigma L4386), 3.574 g HEPES, 100 mg gentamicin sulfate (Sigma G-1264). The pH of the media was adjusted to 7.4 using NaOH.

At day of assay, oocytes were washed twice with room temp OR2 buffer (82.5 mM NaCl, 2.5 mM KCl, 1 mM CaCl_2_, 1 mM MgCl_2_, 1 mM Na_2_HPO_4_, and 15 mM HEPES pH 7.4). Substrates were added to oocytes as a 10x stock. Incubation time was 20 minutes unless specified in figures. Transport was terminated by putting the plate on ice. Oocytes were immediately transferred into ice-cold OR2 buffer, and washed four more times with ice-cold OR2 buffer. Individual oocytes were lysed in acetonitrile:water (90:10, v/v). Homogenates were centrifuged twice at 21,130 × g for 30 min at 4 °C, and supernatants were used for LC-MS analysis.

### Mammalian cell-based transport assay

HCT116 MCT6-KO2 cells stably expressing doxycycline-inducible WT or mutant mouse MCT6 were seeded at 1 million cells per well in six-well plates. The following day, media were replaced with fresh media containing either 1 µg/mL doxycycline (to induce MCT6 expression, OE) or vehicle (0.01% ethanol; uninduced KO control). After 36 hours, cells were washed twice with room-temperature HBSS and then incubated with 1 mL of HBSS containing 1 mM (unless otherwise noted in the figure) deuterated Lac-Phe (D_3_-Lac-Phe) for 10 min at 37°C. To determine the Na⁺ and Cl⁻ dependence of Lac-Phe transport, HBSS was prepared with Na⁺ replaced by NMDG^+^, Cl⁻ replaced by gluconate, or both ions replaced simultaneously with mannitol matching the osmolarity. Transport assays were performed as described above, with a 2 min incubation. For pH-dependence assay, HBSS was buffered with 10 mM MES (pH 5.5, 6.0, 6.5, or 7.0) or 10 mM HEPES (pH 7.5 or 8.0). Transport assays were performed as described above, with a 10 min incubation. For the membrane-potential assay, buffers contained 140 mM NaCl or KCl, 10 mM glucose, 20 mM MES (pH 6.0) or HEPES (pH 7.4), 1.8 mM CaCl₂, and 1 mM MgCl₂, with valinomycin and nigericin each added to a final concentration of 10 µM to dissipate membrane potential and pH gradients, respectively. Cells were preincubated in the appropriate buffer for 15 min, followed by a 2 min transport assay as described above.

Transport was terminated by immediately placing the plate on ice and washing cells twice with ice-cold saline. Cells were lysed in 500 µL of acetonitrile:water (90:10, v/v), and lysates were centrifuged at 21,130 × g for 20 min at 4°C. The supernatant was collected for LC-MS analysis.

### Molecular docking

Molecular docking of Lac-Phe onto mouse MCT6 was performed using DiffDock. The AlphaFold-predicted structure of murine MCT6 (AF-G5E8K6-F1) was used and prepared as a PDB file. The substrate Lac-Phe was prepared as an SDF file. Predicted docking poses were evaluated in PyMOL, and residues predicted to line the active site were selected for subsequent mutagenesis.

### Binding pocket comparison

Putative substrate-binding pockets of MCT6, ABCC5, SLC17A1, and SLC17A3 were compared using SiteMap (Schrödinger Maestro). The same AlphaFold-predicted MCT6 structure used for docking was analyzed. ABCC5 was analyzed using the inward-open cryo-EM structure (PDB 8WI3). SLC17A1 (UniProt Q14916) and SLC17A3 (UniProt O00476) were analyzed using their corresponding AlphaFold-predicted structures.

### MCT6 mutagenesis

Mouse MCT6 cDNA was amplified from the plasmid (Origene, MR221292). A Q5 Site-Directed Mutagenesis Kit (NEB, E0554S) was used to introduce mutations in amino acid residues predicted to play a role in interacting with Lac-Phe on the pCMV vector. WT MCT6 and mutants were subsequently cloned into pCW57.1_TStrp_HA_Ct and transduced into HCT116 MCT6-KO2 cells to allow doxycycline-inducible expression. The sequences were verified through whole plasmid sequencing by Plasmidsaurus.

### Immunocytochemistry and confocal image acquisition

Cells were fixed with 4% paraformaldehyde and 4% sucrose in PBS for 10 min at room temperature, followed by three washes with PBS. Cells were then permeabilized and blocked in PBS containing 0.2% Triton X-100 and 5% goat serum for 30 min at room temperature. Primary antibody incubation was performed with rabbit anti-HA antibody for 60 min at room temperature. After three washes with PBS, cells were incubated with Alexa Fluor 546-conjugated goat anti-rabbit secondary antibody (1:1,000) for 60 min at room temperature. Following three additional PBS washes, coverslips were mounted onto UltraClear microscope slides (Denville Scientific) using Fluoromount-G mounting medium containing DAPI (Invitrogen, 00-4959-52).

Serial confocal z-stack images were acquired using a Nikon A1RSi confocal microscope equipped with a 60× oil-immersion objective. Images were captured and analyzed using NIS-Elements AR software. Laser intensities and acquisition settings for each channel were optimized using appropriate LUT settings and kept consistent across all experimental replicates. All staining procedures and imaging parameters were maintained uniformly throughout experiments.

### Metabolite extraction from mouse plasma

Blood was collected from mice through a submandibular bleeding into lithium heparin tubes (BD, 365985) and immediately kept on ice. The blood was then centrifuged at 4 °C for 5 min at 2,348 x g. Metabolites for LC-MS analysis were extracted by adding 150 µL of 2:1 mixture (v/v) of acetonitrile:methanol to 50 µL plasma. The mixture was then mixed by vortex and centrifuged at 4°C for 10 min at 21,130 x g. The supernatant was transferred to a LC-MS vial. To enrich lactoyl amino acids, 50 µL plasma (after adding HCl, final 100 mM HCl) were mixed with 600 µL ethyl acetate, with 540 µL top layer dried down under nitrogen. The mixture was resuspended in 100 µL of acetonitrile:water (90:10, v/v). For non-terminal bleeding with multiple time points, 30 µL plasma (after adding HCl to a final concentration of 100 mM) were mixed with 600 µL ethyl acetate, with 540 µL top layer dried down under nitrogen. The mixture was resuspended in 50 µL acetonitrile:water (90:10, v/v). For both protocols, the mixture was centrifuged at 4 °C for 10 min at 21,130 x g and the supernatant was transferred to an LC-MS vial.

### Metabolite extraction from mouse tissues

Mouse tissues were dissected and immediately frozen in liquid nitrogen. Around 100 mg tissue was homogenized in 500 µL HPLC-grade water with a Benchmark Beadblaster Homogenizer at 4°C. The homogenate was centrifuged at 4°C for 10 minutes at 21,130 x g. For metabolome analysis, 50 µL of supernatant was added to 150 µL of a 2:1 (v/v) mixture of acetonitrile:methanol. The mixture was centrifuged at 4°C for 10 minutes at 21,130 x g and the supernatant was transferred to a LC-MS vial. For lactoyl amino acid analysis, 300 µL of the supernatant was acidified with 20 µL 1M HCl, then extracted with 600 µL ethyl acetate. The mixture was mixed by vortexing for 30 seconds. After a centrifugation at 4°C for 1 min at 21,130 x g, 300 µL of the top layer was dried down and reconstituted with 200 µL acetonitrile:water (90:10, v/v). The mixture was centrifuged at 4°C for 10 min at 21,130 x g and the supernatant was transferred to an LC-MS vial. A few samples (3 intestine tissue samples) were heavily emulsified with fat and did not yield enough ethyl acetate layer for later analysis and therefore were excluded from further analysis. Metabolite levels were normalized to tissue weight.

### LC-MS analysis

Untargeted metabolomics measurements were performed using an Agilent 6520, 6530 or 6545 Quadrupole time-of-flight LC-MS instrument. MS analysis was performed using electrospray ionization (ESI). The dual ESI source parameters were set as follows: the gas temperature was set at 250 °C with a drying gas flow of 12 L/min and the nebulizer pressure at 35 psi; the capillary voltage was set to 3,500 V; and the fragmentor voltage set to 100 V. Separation of metabolites was conducted using either a Luna 5 μm NH_2_ 100 Å LC column (Phenomenex, 00B-4378-E0) or a Poroshell 120 HILIC-Z 2.1 x 150 mm, 2.7 µm (Agilent, 683775-924) with normal phase chromatography. For NH2 column, mobile phases were as follows: buffer A, 95:5 water:acetonitrile with 0.2% ammonium hydroxide and 10 mM ammonium acetate; buffer B, acetonitrile. The LC gradient started at 100% buffer B with a flow rate of 0.7 mL/min from 0 to 2 min. The gradient was then linearly increased to 50% buffer A and 50% buffer B at a flow rate of 0.7 mL/min from 2 to 20 min. From 20 to 25 min, the gradient was maintained at 50% buffer A and 50% buffer B at a flow rate of 0.7 mL/min. For HILIC-Z column, mobile phases were as follows: buffer A, water with 10 mM ammonium acetate, pH 9.0; buffer B, 90:10 acetonitrile:water with 10 mM ammonium acetate, pH 9.0. The LC gradient started at 100% buffer B from 0 to 2 min. The gradient was then linearly increased to 30% buffer A and 70% buffer B from 2 to 12 min. The gradient was linearly increased to 60% buffer A and 40% buffer B from 12 to 14 min, and kept at this ratio for another minute. Then the gradient returned to 100% B for 5 minutes to re-equilibrate the column. The flow rate was kept constant at 0.25 mL/min.

### Untargeted metabolomics identification of MCT6 substrates

Differential peak identification between different groups was first performed with XCMS (Version 3.5.1) with the following feature detection parameters: 30 ppm, signal/noise threshold 20, mzdiff 0.1. A meta-analysis was performed to identify features commonly shared across different cell lines in the overexpression experiments, or across independent pooled knockout lines in the KO experiments. The following parameters were used: filtering, pval 0.05 (after BH-FDR correction), fc 1.5, maxint 20000; Alignment, mztol 0.01, rttol 60. A full list of XCMS results was attached as **Table S2**.

### Effect of metformin treatment on food intake and body weight

Male WT and MCT6-KO mice were generated by heterozygous breeding crosses. At 4-6 weeks, mice were switched from chow diet to HFD. At 10-14 weeks, mice were single housed and mock gavaged with saline 3-5 days till body weight stabilized to reduce stress-induced loss of food intake and body weight. Male WT and MCT6-KO were body weight matched and received metformin (300 mg/kg unless specified, p.o.). Body weight and food intake were measured daily. For the month-long treatment, mice were 17-18 weeks old, on HFD for 11-12 weeks. Then mock gavaged for 3 days before the treatment with metformin. Body weight and food intake were measured daily. For the experiment where metformin and Lac-Phe were co-administered, 15-22 week old male mice which have been on HFD for 11-12 weeks, were single housed and mock gavaged with i.p. injections daily before the experiment. For the gut-specific MCT6-KO mice, *Vil1^Cre+/-;^ Slc16a5^fl/fl^* mice and no Cre littermates were 14-18 weeks old, and were on HFD for 9 weeks.

### Hormone measurements after metformin treatment

Male WT and MCT6-KO DIO mice (12-14 weeks old), or male gut-specific MCT6-KO and controls (10-13 weeks old) were treated with one single dose of metformin (300 mg/kg, p.o.) and blood was taken 4 hours after the treatment. Levels of hormones were measured with commercial ELISA kits following the manufacturers’ instructions. Information about the ELISA kits is provided below: Mouse PYY (Crystal Chem, 81501), Multi Species GLP-1 (Sigma, EZGLP1T), Mouse/Rat GDF15 (R&D, MGD150).

### Glucose tolerance test (GTT)

Male WT and MCT6-KO DIO mice (12-14 weeks old) were single housed and fasted for 6 hours. The mice were then gavaged with metformin (300 mg/kg, p.o.). 30 minutes later, the mice were injected with glucose (1 g/kg, i.p.). Blood glucose was measured at 0, 20, 40, 60, 90, 120 minutes after the glucose injection.

### Indirect calorimetry and physiological measurements

Male MCT6-KO and WT mice (25-30 weeks old, after 12-16 weeks of high fat diet) were used. Metabolic parameters including oxygen consumption, carbon dioxide production, respiratory exchange ratio (RER), and ambulatory movement of mice were measured using the environment-controlled home-cage CLAMS system (Columbus Instruments) at the Stanford Diabetes Center.

### Mouse running protocols

For mouse running studies a 6 lane Columbus Instruments animal treadmill (product 1055-SRM-D65) was used. For the moderate treadmill running protocol: mice were first acclimated to the treadmill for 5 minutes before the running protocol began. The treadmill running was performed at a constant 5° incline and began at a speed of 6 m/min. Speed was increased by 2 m/min every 5 minutes. For biochemical analyses on blood plasma metabolites, mice were stopped at 30 minutes and blood was taken at time points indicated. For chronic running experiments, mice were exercised 5 days per week (Monday-Friday) and kept on high fat diet (60% kcal from fat). Treadmill was set as above with a maximum of 30 m/min. Mice were stopped upon reaching exhaustion and running times were normalized between the two genotypes. For acute exercise to exhaustion (maximum effort, high-incline treadmill running) study, treadmill running began at a speed of 7.5 m/min and a 4° incline. Every 3 min, the speed and incline were increased by 2.5 m/min and 2°, respectively. Once a speed of 40 m/min and an incline of 30° were reached, both parameters were kept constant until mice reached exhaustion. Exhaustion was defined as remaining on the shocker at the rear of the treadmill for longer than 5 seconds.

### Statistics

All data are expressed as mean ± SEM. *P*-values and statistical methods are provided in figure legends.

## Resource availability

### Lead contact

Further information and requests for resources and reagents should be directed to and will be fulfilled by the lead contact, Jonathan Long.

### Materials availability

Chemical compounds generated in this study can be directly requested by email to the lead contact and are available without restrictions.

### Data and code availability

All data reported in this paper will be shared by the lead contact upon request. This paper does not report original code. Any additional information requested to re-analyze the data reported in this paper is available from the lead contact upon request.

## Acknowledgements

We thank Dr. Zhiwen Liao of the Goodman lab at Stanford for her exceptional technical work enabling oocyte expression studies. We thank Dr. Hong Zeng from the transgenic mouse facility for help in generating the conditional allele of *Slc16a5*. We thank Dr. Liang Feng and members of the Long laboratory for helpful discussions. We thank B. Gallardo for assistance in laboratory operations. This work was supported by the NIH (R01DK124265 and DP1DK130641 to J.Z.L., K99DK141966 to S.X., R21DC021031 and R01GM139936 to J.A.S., R21EY036218 to M.D.P.), the National Science Foundation (CAREER Award to J.A.S.), the Stanford Diabetes Research Center (P30DK116074 to J.Z.L.), the Phil & Penny Knight Initiative for Brain Resilience at the Wu Tsai Neurosciences Institute (research grant to J.Z.L.), the Ono Pharma Foundation (research grant to JZL), the Weill Cancer Hub West (research grant to J.Z.L.), and the Stanford Wu Tsai Human Performance Alliance (research grant to J.Z.L. and postdoctoral fellowship to S.X., M.D.M.G., and X.L.).

## Author contributions

Conceptualization: S.X., J.Z.L., A.S.-H.T. Investigation: S.X., A.S.-H.T., J.S., S.F., X.C., M.D.M.G., K.T., H.C., K.I., X.L., V.L.L., H.T.C., K.I., S.C.R., C.L., X.L., S.H.R., M.D.W.T., A.G., J.T., M.Y., W.W., S.M.H. Writing: S.X., J.Z.L., S.M.H., A.S.-H.T. Resources: J.Z.L., M.R.H., M.D.P., J.A.S. Supervision, J.Z.L.

## Supplementary Figure Legends

**Figure S1.**
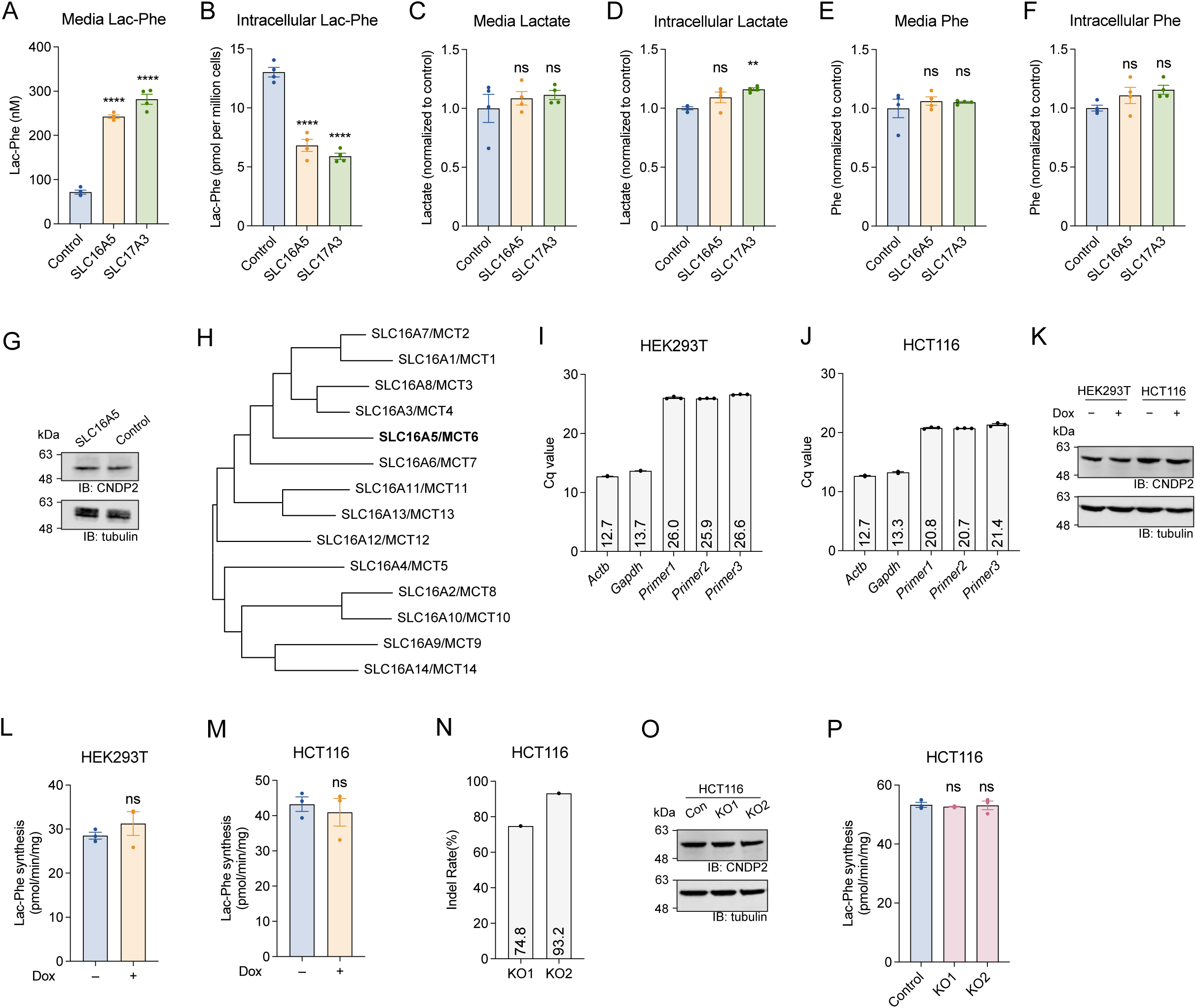
Additional characterization of MCT6 overexpressing or knockout cells. (A,B) Levels of Lac-Phe in conditioned medium (A) or cell lysates (B) of transfected HEK293T cells expressing mCherry (Control), SLC16A5, or SLC17A3. N = 4/group. (C,D) Levels of lactate in conditioned medium (C) or cell lysates (D) of transfected HEK293T cells expressing mCherry (Control), SLC16A5, or SLC17A3. N = 4/group. (E,F) Levels of phenylalanine in conditioned medium (E) or cell lysates (F) of transfected HEK293T cells expressing mCherry (Control), SLC16A5, or SLC17A3. N = 4/group. (G) Anti-CNDP2 (top) and anti-tubulin (bottom) blotting of HEK293T cells transfected with SLC16A5 or mCherry (Control). (H) Phylogenetic tree of the SLC16/MCT transporter family. The phylogeny was constructed using the Maximum Likelihood method using protein sequences. (I) Expression levels of *Actb*, *Gapdh*, and *Slc16a5* mRNA in HEK293T (I) and HCT116 (J) cells. N = 3 technical replicates. (K) Anti-CNDP2 (top) and anti-tubulin (bottom) blotting of HEK293T and HCT116 cells with or without doxycycline-inducible MCT6 expression. (L,M) Lac-Phe synthesis activity of cell lysates of HEK293T (L) and HCT116 (M) cells with or without doxycycline-inducible MCT6 expression. N = 3/group. (N) Indel rate of HCT116 KO1 and KO2 cells. (O) Anti-CNDP2 (top) and anti-tubulin (bottom) blotting of HCT116 cells with corresponding genotypes. (P) Lac-Phe synthesis activity of cell lysates of HCT116 cells with corresponding genotypes. N = 3/group. Data are shown as mean ± SEM. ** *p* < 0.01, **** *p* < 0.0001. In (A-F,P) *p*-values were calculated from one-way ANOVA with post hoc Dunnett’s multiple comparisons test. In (L,M), *p*-values were calculated with two-tailed unpaired t tests.

**Figure S2.**
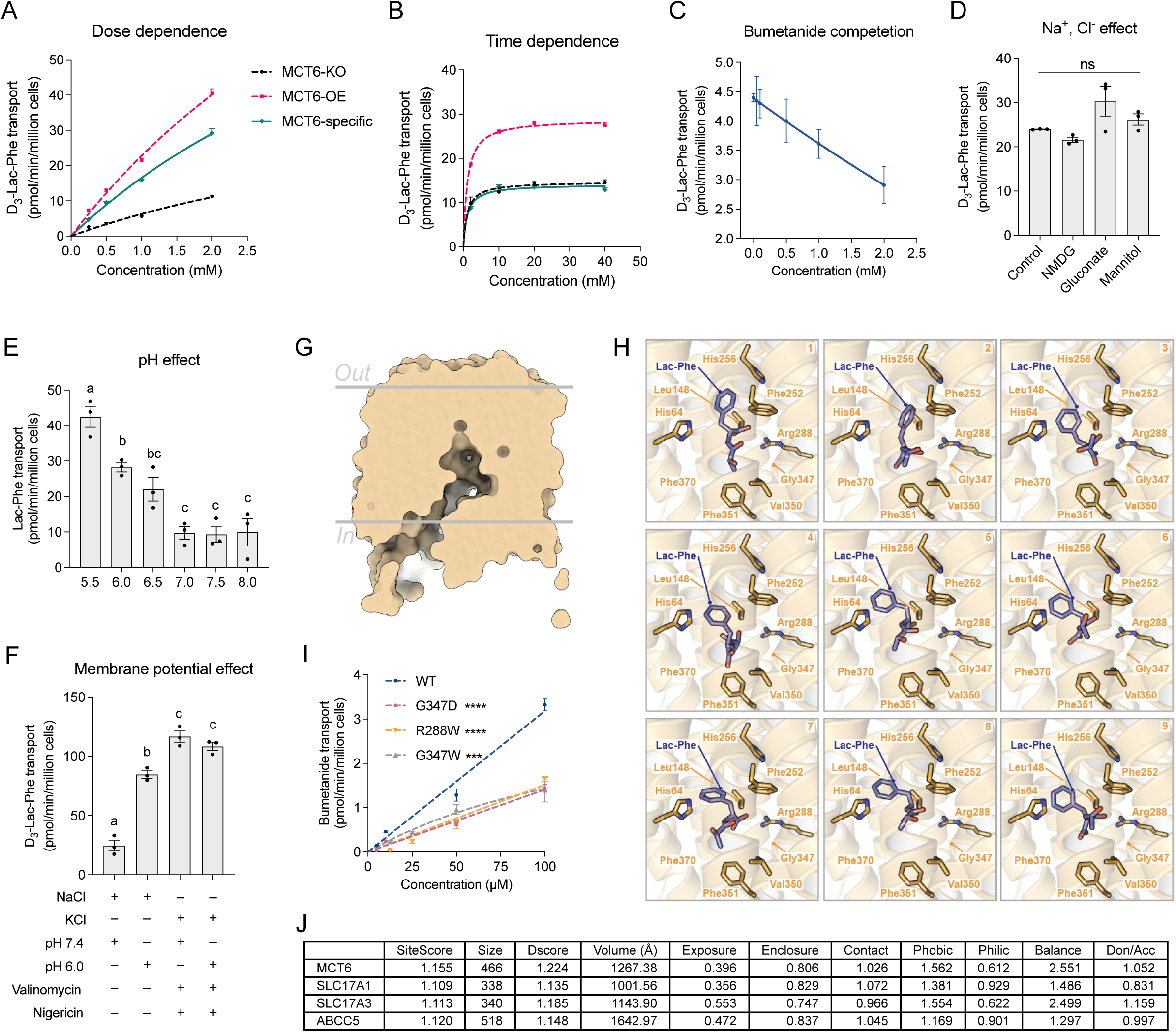
Additional characterization of MCT6-mediated Lac-Phe transport. (A) D_3_-Lac-Phe transport activity in MCT6-KO cells (black line) or in KO cells reconstituted with MCT6 (pink line) with different D_3_-Lac-Phe concentrations for 10 minute incubation (A) or with different incubation times and 1 mM D_3_-Lac-Phe (B). MCT6-specific transport activity is shown in the teal line. N = 3/time point. (C) MCT6-specific D_3_-Lac-Phe transport activity in a mammalian cell-based transport assay with 1 mM D_3_-Lac-Phe in the presence of various concentrations of bumetanide for 10 minute incubation. N = 3/group. (D) MCT6-specific D_3_-Lac-Phe transport activity in a mammalian cell-based transport assay with 1 mM D_3_-Lac-Phe with standard HBSS (Control), Na^+^-free HBSS (NMDG), Cl^−^-free HBSS (Gluconate), or Na^+^,Cl^−^-free HBSS (Mannitol). N = 3/group. (E) MCT6-specific Lac-Phe transport activity in a mammalian cell-based transport assay with 1 mM Lac-Phe at different pHs for 10 minutes. N = 3/group. (F) MCT6-specific D_3_-Lac-Phe transport activity in a mammalian cell-based transport assay with 1 mM D_3_-Lac-Phe at pH 6.0 or 7.4. Valinomycin and nigericin in a high KCl buffer was used to dissipate both the membrane potential and proton gradient. N = 3/group. (G) Inward-open conformation of mMCT6 based on an AlphaFold-predicted structure. (H) Additional top-ranked molecular docking poses of mMCT6 and Lac-Phe. Individual residues (orange) within 4Å of Lac-Phe (blue) are highlighted. (I) MCT6-specific bumetanide transport activity of WT, G347D, R288W, and G347W MCT6 mutants. N = 3/group. (J) Physiochemical properties of putative substrate-binding cavities in MCT6, SLC17A1, SLC17A3, and ABCC5 using SiteMap. SiteScore, overall composite site quality score; Size, number of site points used to define the pocket; Dscore, druggability score; Volume, physical volume of the identified pocket; Exposure, ratio of the pocket exposed to solvent; Enclosure, ratio of the pocket surrounded by protein structure; Contact, degree of direct van der Waals contact; Phobic, hydrophobic character score of the pocket lining; Philic, hydrophilic character score of the pocket lining; Balance, ratio of Phobic to Philic; Don/Acc, ratio of hydrogen-bond donor character to acceptor character. For (A-F) and (I), MCT6-specific transport activity was determined by subtraction of the background activity in the MCT6-KO cells. Data are shown as mean ± SEM. *** *p* < 0.001, **** *p* < 0.0001. In (D), *p*-values were calculated from one-way ANOVA with post hoc Dunnett’s multiple comparisons test, compared to Control. In (E,F), *p*-values were calculated from one-way ANOVA with post hoc Tukey’s multiple comparisons test, different letters represent *p*-values < 0.05. In (I), *p*-values were calculated from mixed-effects analysis with Tukey’s multiple comparisons tests.

**Figure S3.**
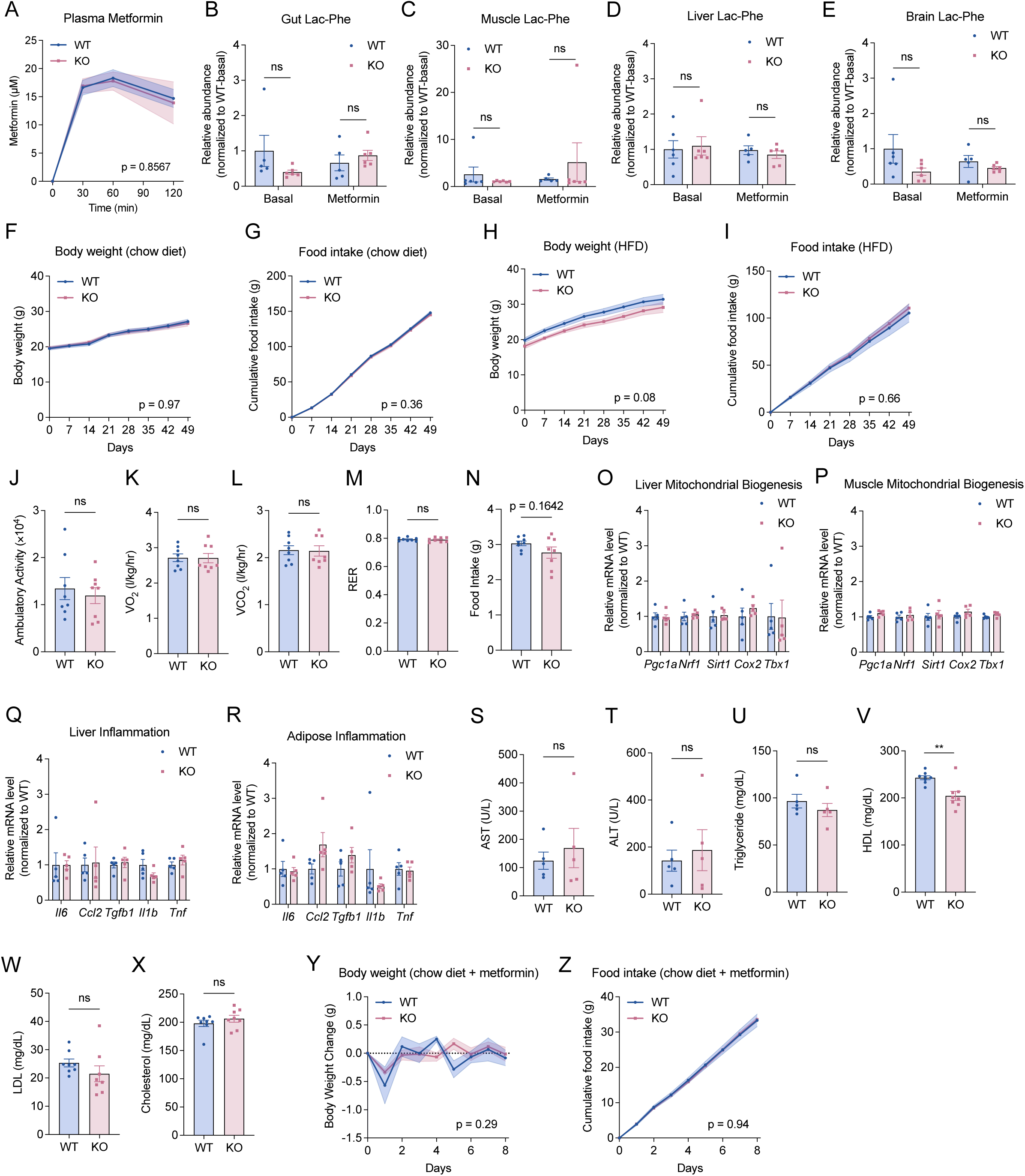
Additional molecular characterization of MCT6-KO mice. (A) Plasma metformin levels in WT and MCT6-KO mice after a single oral administration (300 mg/kg, p.o.) (B-E) Levels of Lac-Phe in the gut (B), muscle (C), liver (D), and brain (E) at both basal and metformin-induced (30 min after a single administration of metformin, 300 mg/kg, p.o.) states in 14-20 week-old female WT and MCT6-KO mice. (F,G) Cumulative change in body weight (F) and food intake (G) in 4-6 week old male WT and MCT6-KO mice on chow diet. N = 12 for WT and N = 7 for KO. (H,I) Cumulative change in body weight (H) and food intake (I) in 4-6 week old male WT and MCT6-KO mice on high fat diet (HFD). N = 6 for WT and N = 10 for KO. (J-N) 24 hour locomotor activity (J), oxygen consumption (K), carbon dioxide production (L), respiratory exchange ratio (RER, M), and food intake (N) of 25-30 week-old male WT or MCT6-KO mice after 12-16 weeks of high fat diet. N = 7-8/group. (O-R) Relative mRNA expression of indicated genes in liver (O,Q), quadriceps muscle (P) or epididymal white adipose tissue (R) from 25-30 week-old male WT and MCT6-KO mice on HFD for 12-16 weeks. N = 5/group. (S-X) Levels of AST (S), ALT (T), triglycerides (U), HDL (V), LDL (W), cholesterol (X) from 25-30 week old male WT and MCT6-KO mice on HFD for 12-16 weeks. N = 5/group. (Y,Z) Cumulative change in body weight (Y) and food intake (Z) in 20-24 week-old male WT and MCT6-KO lean mice following chronic metformin treatment (300 mg/kg daily, p.o.). N = 6/group. Starting body weights were WT: 31.5 ± 1.9 g, KO: 31.6 ± 1.4 g (mean ± SEM). Data are shown as mean ± SEM. ** *p* < 0.01. In (B-E,O-R), *p*-values were calculated with two-way ANOVA with post hoc Sidak’s multiple comparisons test. In (F-I,Y,Z), *p*-values were calculated from two-way ANOVA and reporting the effect of genotype. In (J-N,S-X), *p*-values were calculated from two-tailed unpaired t-test.

**Figure S4.**
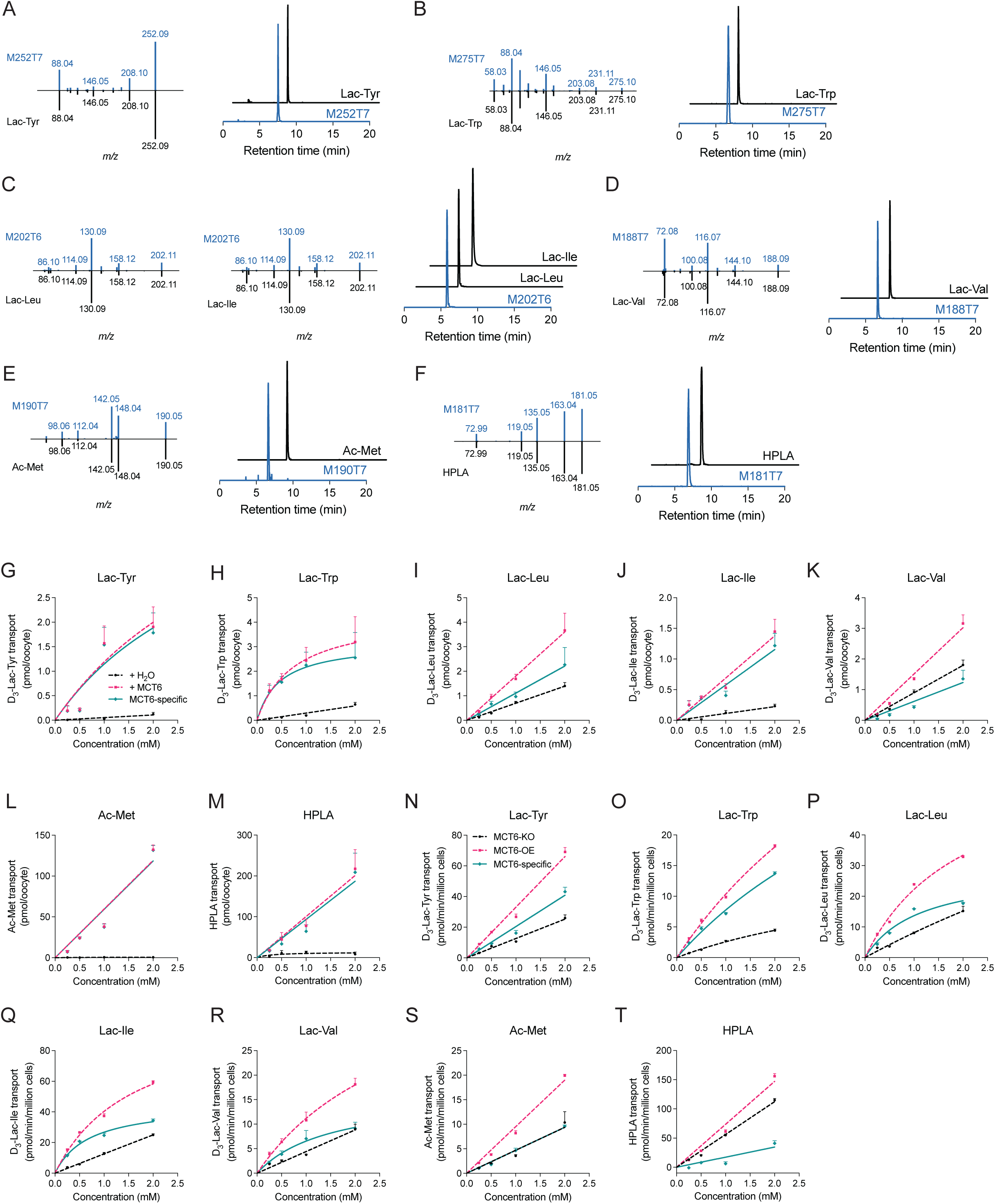
Characterization of additional MCT6 substrates. (A) MS/MS fragmentation of the endogenous *m/z* = 252.09 (left, top) and authentic Lac-Tyr standard (left, bottom) and co-elution of the endogenous peak and synthesized standard (right). (B) MS/MS fragmentation of the endogenous *m/z* = 202.11 (left and middle, top), authentic Lac-Leu (left, bottom) and Lac-Ile (middle, bottom) standard, and co-elution of the endogenous peak and synthesized standards (right). (C) MS/MS fragmentation of the endogenous *m/z* = 188.09 (left, top) and authentic Lac-Val standard (left, bottom) and co-elution of the endogenous peak and synthesized standard (right). (D) MS/MS fragmentation of the endogenous *m/z* = 275.10 (left, top) and authentic Lac-Trp standard (left, bottom) and co-elution of the endogenous peak and synthesized standard (right). (E) MS/MS fragmentation of the endogenous *m/z* = 190.05 (left, top) and authentic Ac-Met standard (left, bottom) and co-elution of the endogenous peak and synthesized standard (right). (F) MS/MS fragmentation of the endogenous *m/z* = 181.05 (left, top) and authentic HPLA standard (left, bottom) and co-elution of the endogenous peak and synthesized standard (right). (G-M) Transport activity of *X. laevis* oocytes with different concentrations of additional MCT6 substrates: Lac-Tyr (G), Lac-Trp (H), Lac-Leu (I), Lac-Ile (J), Lac-Val (K), , Ac-Met (L), HPLA (M). N = 3-5/group (N-T) Transport activity with a mammalian cell-based transport assay with different concentrations of additional MCT6 substrates: Lac-Tyr (N), Lac-Trp (O), Lac-Leu (P), Lac-Ile (Q), Lac-Val (R), Ac-Met (S), HPLA (T). For (G-T), MCT6-specific transport activity was determined by subtraction of the background activity in water-injected oocytes (G-M) or MCT6-KO cells (N-T). N = 3/group. Data are shown as mean ± SEM.

**Figure S5.**
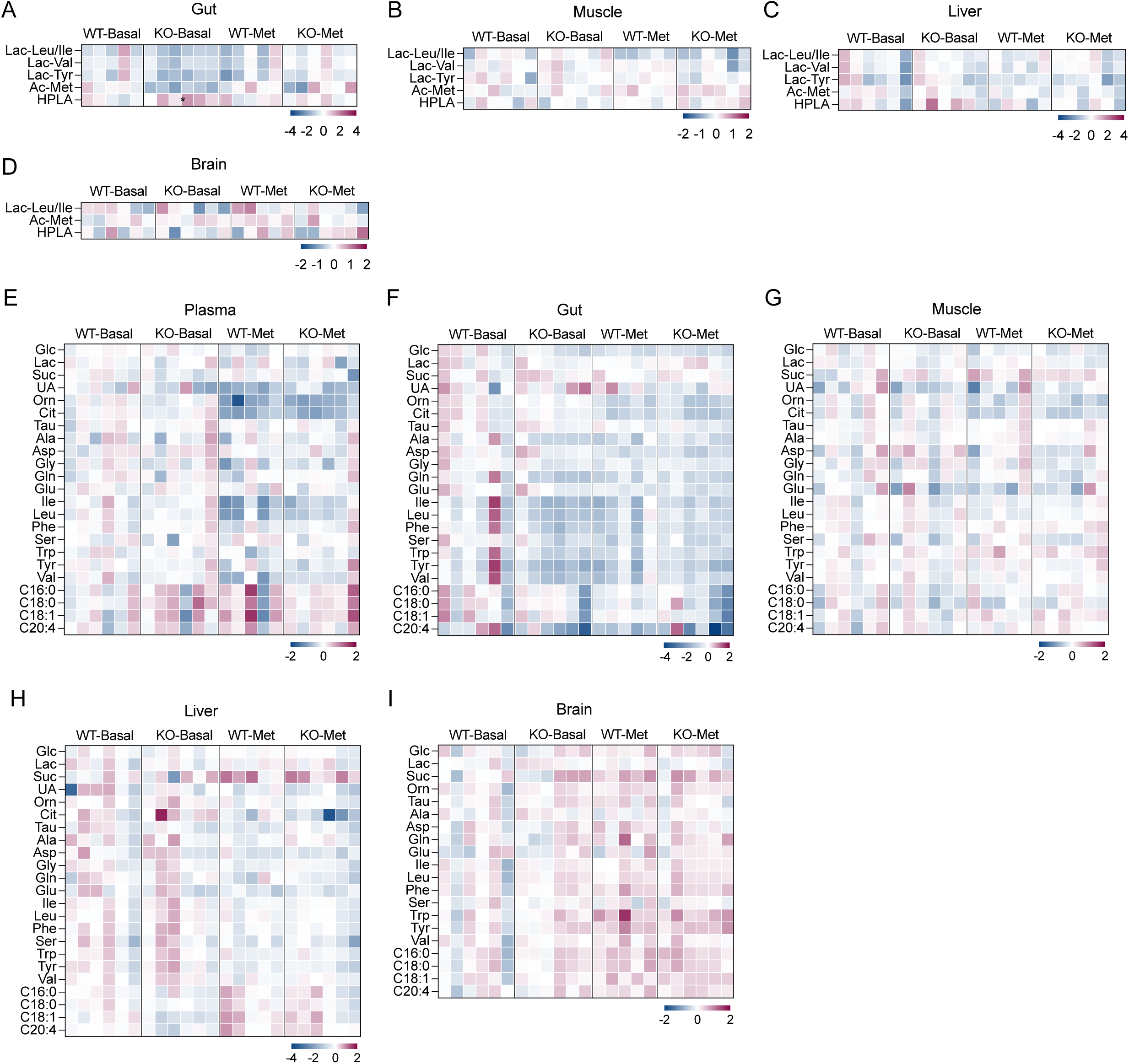
Biochemical characterization of MCT6-KO mice. (A-D) Levels of MCT6 substrates in the gut (A), muscle (B), liver (C), and brain (D) of 14-20 week-old female WT and MCT6-KO mice under basal state or 30 min after a single administration of metformin (300 mg/kg, p.o.). Levels were normalized to WT mice under basal state, with log_2_ transformation. Lac-Trp was below detection limit in all four tissues, Lac-Tyr and Lac-Val were below detection limit in brain tissues. (E-I) Levels of metabolites in the blood plasma (E), gut (F), muscle (G), liver (H), and brain (I) of 14-20 week-old female WT and MCT6-KO mice under basal state or 30 min after a single administration of metformin (300 mg/kg, p.o.). Levels were normalized to WT mice under basal state, with log_2_ transformation. Glc, glucose; Lac, lactate; Suc, succinate; UA, uric acid; Orn, ornithine; Cit, citrulline; Tau, taurine; Ala, alanine; Asp, aspartic acid; Gly, glycine; Gln, glutamine; Glu, glutamic acid; Ile, isoleucine; Leu, leucine; Phe, phenylalanine; Ser, serine; Trp, tryptophan; Tyr, tyrosine; Val, valine; C16:0, palmitate; C18:0, stearate; C18:1, oleate; C20:4, arachidonate. Data are shown as mean ± SEM. * *p* < 0.05. In all panels, p-values were calculated separately for Basal state and Metformin state, from two-way ANOVA with post hoc Šídák’s multiple comparisons test.

**Figure S6.**
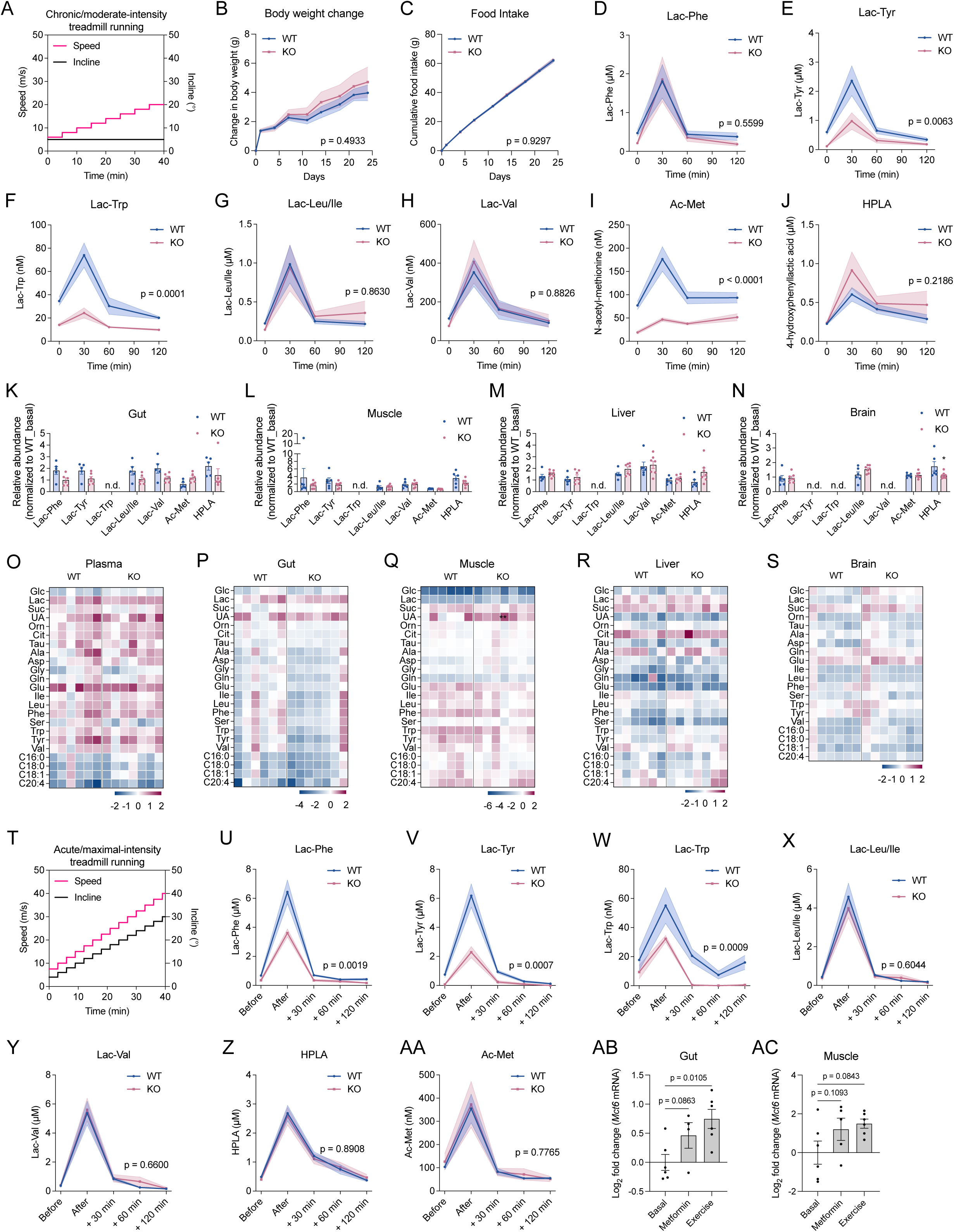
Effect of treadmill exercise in MCT6-KO mice. (A) Schematic of speed and incline in the chronic/moderate intensity running protocol for mice. (B,C) Cumulative body weight change (B) and cumulative food intake (C) in 10-14 week old male WT and MCT6-KO mice following a combined chronic treadmill running and high fat diet feeding protocol. N = 8 for WT and N = 7 for KO. (D-J) Plasma levels of Lac-Phe and additional MCT6 substrates at the indicated time point from 8-12 week old male WT and MCT6-KO mice before (time 0) and after a single bout of moderate intensity treadmill running for 30 min. N = 8 for WT, N = 6 for KO. (K-N) Levels of MCT6 substrates in the gut (K), muscle (L), liver (M), and brain (N) of 14-20 week-old female WT and MCT6-KO mice under basal state or after a single bout of treadmill running for 30 min. Levels were normalized to WT mice under basal state, with log_2_ transformation. N = 6 for WT, N = 7 for KO. (O-S) Levels of metabolites in the blood plasma (O), gut (P), muscle (Q), liver (R), and brain (S) of 14-20 week-old female WT and MCT6-KO mice under basal state or after a single bout of treadmill running for 30 min. Levels were normalized to WT mice under basal state, with log_2_ transformation. N = 6 for WT, N = 7 for KO. Glc, glucose; Lac, lactate; Suc, succinate; UA, uric acid; Orn, ornithine; Cit, citrulline; Tau, taurine; Ala, alanine; Asp, aspartic acid; Gly, glycine; Gln, glutamine; Glu, glutamic acid; Ile, isoleucine; Leu, leucine; Phe, phenylalanine; Ser, serine; Trp, tryptophan; Tyr, tyrosine; Val, valine; C16:0, palmitate; C18:0, stearate; C18:1, oleate; C20:4, arachidonate. (T) Schematic of speed and incline in the acute/maximum intensity running protocol for mice. (U-AA) Plasma levels of Lac-Phe and additional MCT6 substrates at the indicated time point from 11-13 week old male WT and MCT6-KO mice before (time 0) and after a single bout of acute/maximum intensity treadmill running. N = 8 for WT, N = 6 for KO. Running time was 29.65 ± 1.6 min for WT, and 29.76 ± 1.7 min for KO. (AB,AC) Change of *Slc16a5* mRNA levels in the gut (AB) or muscle (AC) 30 min after a single administration of metformin (300 mg/kg, p.o.), or a single bout of moderate intensity treadmill running for 30 min. N = 5-6/group. Data are shown as mean ± SEM. ** *p* < 0.01. In (B-J) and (U-AA), *p*-values were calculated from two-way ANOVA and reporting the effect of genotype. In (K-S), *p*-values were calculated from two-way ANOVA with post hoc Šídák’s multiple comparisons test. In (AB,AC), *p*-values were calculated from one-way ANOVA with Holm-Šídák’s multiple comparisons test.

**Figure S7.**
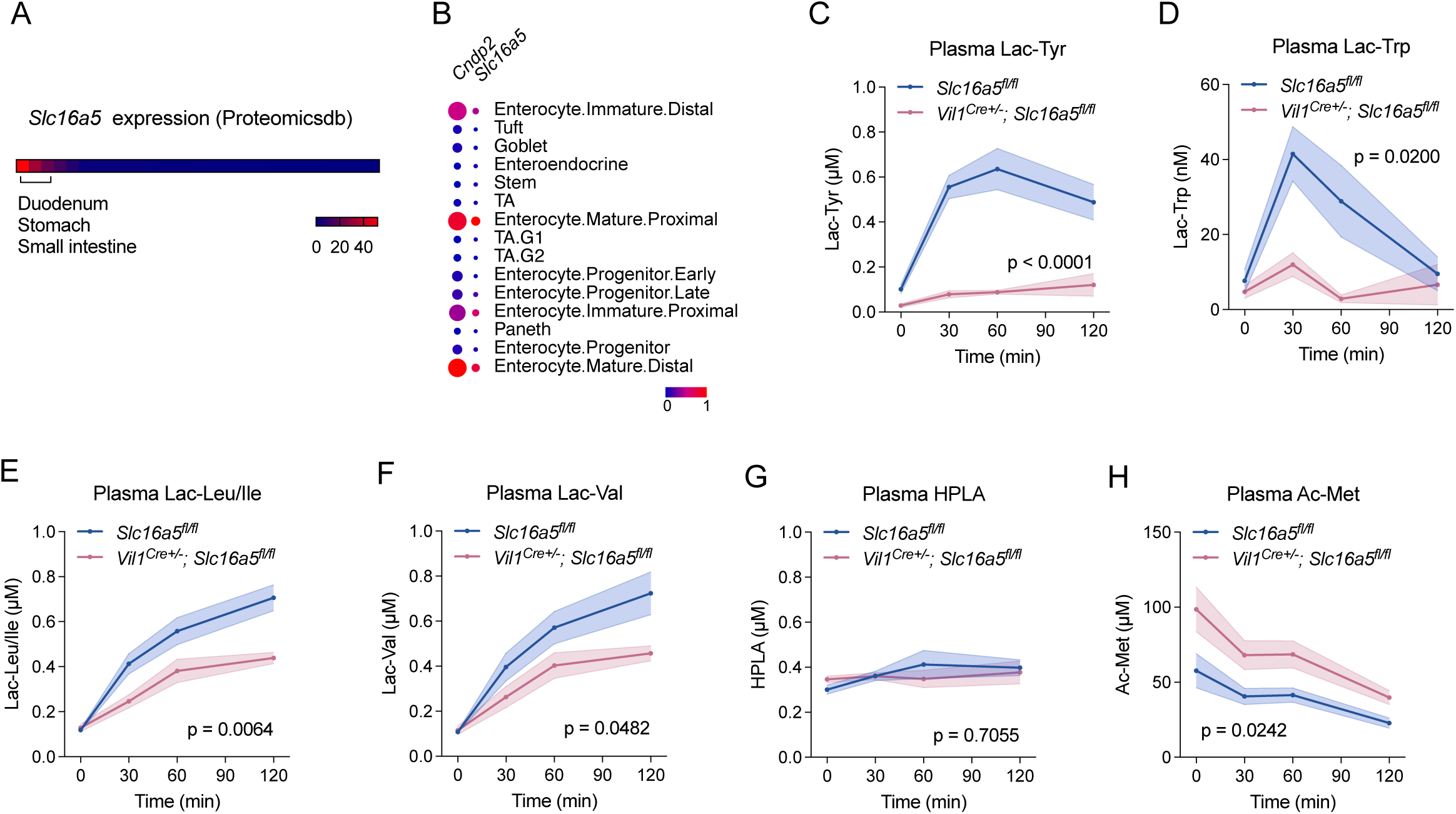
Additional characterization of intestine epithelial-specific MCT6-KO mice. (A) Relative *Slc16a5/Mct6* gene expression (mRNA) level in mouse tissues based on Proteomicsdb dataset. (B) *Slc16a5/Mct6* gene and *Cndp2* gene mRNA gene expression level in single cell mouse small intestine dataset (Haber et al. 2017). (C-H) Plasma levels of additional MCT6 substrates at the indicated time points from 7-8 week-old male *Vil1^Cre+/-^; Slc16a5^fl/fl^* mice and *Slc16a5^fl/fl^* controls before (time 0) and after metformin administration (300mg/kg, p.o.). N = 5-6/group. Data are shown as mean ± SEM. In (C-H), *p*-values were calculated from two-way ANOVA and reporting the effect of genotype.

---

**Table S1. *Cndp2* and transporter co-expression analysis in the gut, related to Fig. 1A**.

**Table S2. Untargeted profiling of MCT6 substrates, related to Fig. 5A**.

